# The genetic and ecological landscape of plasmids in the human gut

**DOI:** 10.1101/2020.11.01.361691

**Authors:** Michael K. Yu, Emily C. Fogarty, A. Murat Eren

## Abstract

Despite their prevalence and impact on microbial lifestyles, ecological and evolutionary insights into naturally occurring plasmids are far from complete. Here we developed a machine learning model, PlasX, which identified 68,350 non-redundant plasmids across human gut metagenomes, and we organized them into 1,169 evolutionarily cohesive ‘plasmid systems’ using our sequence containment-aware network partitioning algorithm, MobMess. Similar to microbial taxa, individuals from the same country tend to cluster together based on their plasmid diversity. However, we found no correlation between plasmid diversity and bacterial taxonomy. Individual plasmids were often country-specific, yet most plasmid systems spanned across geographically distinct human populations, revealing cargo genes that likely respond to environmental selection. Our study introduces powerful tools to recognize and organize plasmids, uncovers their tremendous diversity and intricate ecological and evolutionary patterns in naturally occurring habitats, and demonstrates that plasmids represent a dimension of ecosystems that is not explained by microbial taxonomy alone.

## Main

As a class of mobile genetic elements^1^, plasmids can occur in cells from all domains of life^2,3^, typically as extrachromosomal and circular DNA. Plasmids replicate semi-independently of their hosts, and often transfer between cells as a mechanism of horizontal gene transfer^4–8^. A hallmark of plasmids is their remarkably diverse capacity to impact their microbial hosts through fitness-determining functions they carry^4–6^, such as antibiotic resistance genes^9,10^ and virulence factors^11,12^. Plasmids also exhibit many interesting genetic properties, such as frequent recombination, which can result in plasmids sharing recurrent “backbone” sequences but differing in their cargo genes^13–15^. These backbone sequences often encode for core replication and transfer machinery^13,14,16–18^ that determine the set of compatible hosts they can inhabit^18,19^ as well as regulate their copy number in a specific host^20^. Experiments in model systems and organisms in culture have revealed the critical impact of plasmids in microbial phenotypes and survival especially for pathogens with medical significance. Yet, our understanding of the diversity, ecology, and genetic architecture of naturally occurring plasmids are far from complete.

Recent advances in metagenomics offer unprecedented access to the entire DNA content of an environment without the need for cultivation^21^. In particular, metagenomic assembly and binning strategies have enabled the reconstruction and characterization of microbial genomes de novo^22^, including those in the human gut^23^ where microbes have been associated with health and disease states^24,25^. Metagenomic approaches have also been applied to study plasmid content^26^, but such applications have been limited to shotgun sequencing of plasmid-enriched samples^27–29^ or to surveying only a handful of metagenomes at a time^30–32^. Over the past decade, the number of publicly available metagenomes has rapidly increased, creating an opportunity to conduct large-scale studies to characterize the diversity of naturally occurring plasmids in complex ecosystems.

Several computational strategies have been developed to identify plasmids in sequence collections^17,33,34^. Yet, the ability to distinguish plasmids from bacterial chromosomes or from other mobile genetic elements such as viruses via computational strategies remains a challenge^35^. Popular plasmid prediction strategies rely on k-mer patterns learned from reference plasmid sequences^30,31,36^, exploit known functions such as replication or conjugation genes^37–39^, or use a combination of these features^34^. While these features can help identify plasmids similar to those in public databases, they are of limited utility to recognize novel plasmids. Other approaches focus on circularity of sequences during (meta)genomic assembly^32,40,41^; however, this strategy overlooks plasmids that are linear, integrated, or found as assembly fragments, and may confuse other types of circular mobile elements for plasmids.

Here, we present PlasX, a machine learning approach to identify plasmids in complex microbial ecosystems, and MobMess, a robust network partitioning algorithm to gain insights into plasmid evolution at scale. Using PlasX we identified a collection of 68,350 non-redundant plasmids in the human gut microbiome that were more genetically diverse than reference plasmids and substantially more prevalent across global human populations. We then used MobMess to organize predicted plasmids into ‘plasmid systems’ based on shared backbone sequences, which provided us with an evolutionary framework to investigate plasmid cargo gene content as a function of environmental pressures.

## Results

### A plasmid classification system based on *de novo* gene families

To train our machine learning model, we first compiled a reference set of 16,827 plasmids and 14,367 chromosomal sequences from public databases (Figure 1A, Table S1), in which we identified 51.2 million open reading frames. We were able to assign a function to 71% of plasmid genes using the Cluster of Orthologous Genes (COG)^42^ and/or Pfam^43^ databases. In parallel we clustered all genes into 1,090,132 *de novo* gene families, which accounted for 95% of all plasmid genes (Figures 1B and S1A) and constituted the primary data for the training of PlasX (Figures 1, S1B, and S1C; Supplementary Information).

**Figure 1.**
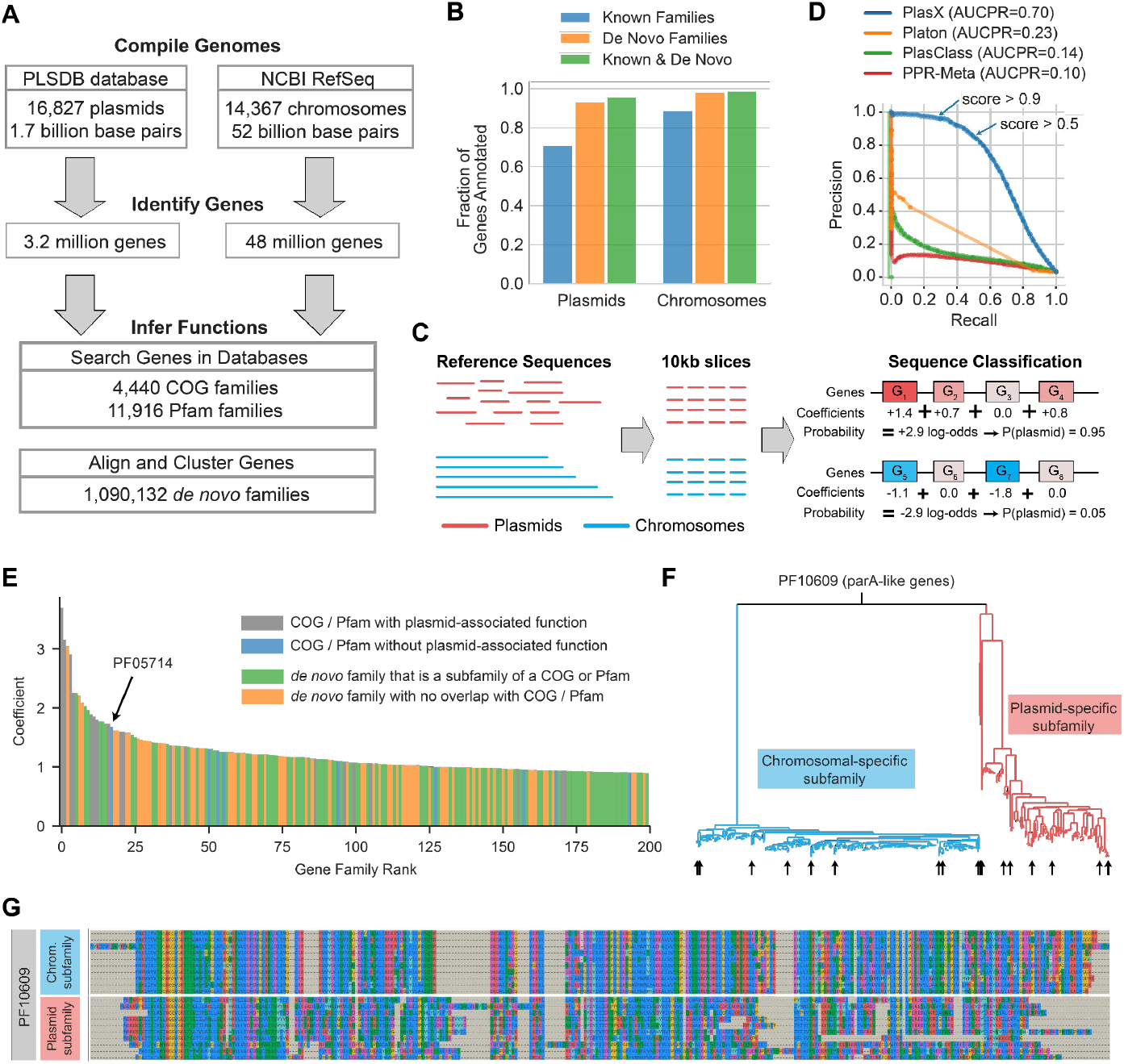
A machine learning model for classifying plasmids. **(A)** Our pangenomics workflow to characterize gene functions in a reference set of plasmids and chromosomes. **(B)** The fraction of all plasmids or all chromosomal genes that are annotated by using known families (blue), *de novo* families (orange), or a combination of both (green). **(C)** Training of PlasX. Reference sequences are sliced into 10 kbp windows and then prediction scores are made by a logistic regression that sums the contributions of gene families within a sequence. **(D)** Precision-recall curves comparing PlasX, Platon, PlasClass, and PPR-Meta. Except for PPR-Meta, every method was trained and evaluated using 4-fold cross-validation and an informed split. AUCPR was calculated using sequence weights for normalization. The arrows indicate the performance of PlasX using a score threshold of either >0.5 or >0.9. **(E)** The 200 gene families with the highest PlasX coefficients and thus most important for identifying plasmids. Gene families are ranked by their coefficient. **(F)** Maximum-likelihood phylogenetic tree of genes that are in PF10609 and also in either the plasmid-specific *de novo* subfamily mmseqs_5_1535552 (red) or chromosome-specific *de novo* subfamily mmseqs_70_40217271 (blue). **(G)** Sequence alignment of 10 representative genes from each subfamily (arrows in F).

PlasX is a logistic regression which assigns a positive or negative coefficient to gene families that are likely to originate from sequences that are of plasmid or non-plasmid origin (Figure 1C). The algorithm predicts whether a given sequence is a plasmid or not by considering the coefficients of all gene families in the sequence, and assigns a score to each prediction that ranges between 0 to 1, where a prediction score over 0.5 suggests that the sequence is more likely to be a plasmid than not. To benchmark PlasX we compared its performance to three state-of-the-art algorithms, PlasClass^31^, PPR-Meta^44^, and Platon^45^ using a cross-validation strategy in which we trained models on a subset of non-redundant reference sequences and evaluated their performance on the remaining sequences that were not used for training (Supplementary Information). PlasX achieved the highest area under the precision-recall curve (AUCPR=0.70) with a substantial improvement compared to the next best method (Platon, AUCPR=0.23) (Figure 1D). We then conducted a test using the 21,012 new plasmids that were recently included in a large plasmid database, PLSDB^46^. PlasX correctly identified 81.5% (17,128) of these new sequences as plasmids, while Platon, the next best method in our cross-validation tests, only predicted 37.4% (7,860) of them as plasmids in its most sensitive mode (Table S2). We further evaluated the performance of PlasX and other methods on a recently characterized novel plasmid of Wolbachia, pWCP^47^. PlasX was able to predict pWCP as a plasmid (score = 0.73), while none of the other methods in our tests recognized pWCP as a plasmid (Table S3). Finally, as an additional benchmarking step to determine PlasX’s ability to distinguish plasmids from other mobile genetic elements, we ran PlasX on all ICE sequences from the ICEberg database^48^ (n=552) and all prophage sequences from the NCBI viral database (n=445). PlasX correctly classified 92.2% of ICEs as not plasmids (Table S4), and 93.2% of NCBI viral database as not plasmids (Table S5). Platon could also distinguish prophages from plasmids with a high accuracy of 99.6%, but its classification accuracy was much lower compared to PlasX’s for ICEs, as Platon classified 37.1% of ICEs as plasmids.

The improved efficiency of PlasX comes from its reliance on de novo gene families rather than sequence features or gene functions alone. Indeed, genes in microbial sequence collections often cannot be annotated to a known function, and a known function often groups together genes with large sequence differences. By partitioning genes into homologous groups, de novo gene families both increase the fraction of input data usable for training and increase the resolution of the resulting units that improve training and prediction. For instance, the Pfam PF10609 is a broad family of genes related to *parA*, a gene that drives the partitioning of not only chromosomes^49^ but also plasmids^50^ during cell division. As genes that resolve to this function are found on 35% of plasmids and 95% of chromosomes, it has no ability to distinguish plasmids and chromosomes (coefficient of -0.023). However, PF10609 in our dataset could be subdivided into two de novo gene families – a plasmid specific one (coefficient +0.455), and a chromosome-specific one (coefficient -0.198). Indeed, the gene sequences in the two de novo gene families of PF10609 show a clear divergence of plasmids and chromosomes into monophyletic groups (Figures 1F, 1G). Furthermore, 35.5% (398,174) of *de novo* gene families did not resolve to a known function, even though many of them had highly positive coefficients. In fact, 12,076 of gene families with unknown functions had coefficients over 0.1, and of the 200 gene families with the highest coefficients, 129 did not have any functional annotation. Our survey of gene families (Table S6) highlights the difficulty of using functional annotations alone to infer the importance of a gene family in plasmid recognition. While some gene families included keywords such as “plasmid”, “replication”, or “conjugation” in their functional descriptions to offer naive confirmations for plasmid relevance, most others were difficult to immediately associate with plasmids. For instance, the gene family with the 17th highest PlasX coefficient of 1.678 in our list was a family of lipoproteins (PF05714). Its annotation, “*Borrelia burgdorferi virulent strain associated lipoprotein*” does not explicitly associate it with plasmids. Yet, it occurred in 168 plasmids and only 2 chromosomes in our training data and has been studied previously for conferring virulence in plasmids^51,52^, suggesting that PlasX-assigned coefficients offer an effective means to identify key gene families to recognize plasmids.

Overall, these results show that PlasX performs better than state-of-the-art plasmid prediction approaches in cross-validation tests, is able to recognize novel plasmids that are not present in existing databases, and is less likely to confuse other mobile genetic elements with plasmids.

### PlasX unveils a large database of new plasmids from the human gut microbiome

Next, we applied PlasX to survey naturally occurring plasmids in the human gut microbiome, an environment which harbors a diverse range of microbes and mobile genetic elements^53^. For this, we assembled 36 million contigs from 1,782 human gut metagenomes, spanning culturally and geographically distinct human populations (Table S7). Running PlasX on these data resulted in a total of 226,194 predicted plasmids with a score above 0.5 (Figures 2A and S2A, Table S8). Our predictions spanned a wide range of lengths, including 135 sequences that were longer than 100 kbp, but they were generally shorter than reference plasmids with a median length of 2.6 kbp versus 53.3 kbp, respectively (Figure S2B). Part of this discrepancy is most likely due to the fragmented nature of assembled sequences from metagenomes. Indeed, the median length of the entire set of contigs was 2.1 kbp, and only 50,310 (0.14%) contigs were longer than 100 kbp. To minimize the impact of assembly fragments in our results, we removed predictions that did not appear to be circular and, at the same time, appeared to be fragments of more complete predictions in our collection. This filter left us with 100,719 predictions for downstream analyses (see Methods, Figure S3). Although this filtered set inevitably contains fragmented plasmids due to the nature of the input data, hereafter we refer to them as ‘plasmids’ for practical reasons.

**Figure 2.**
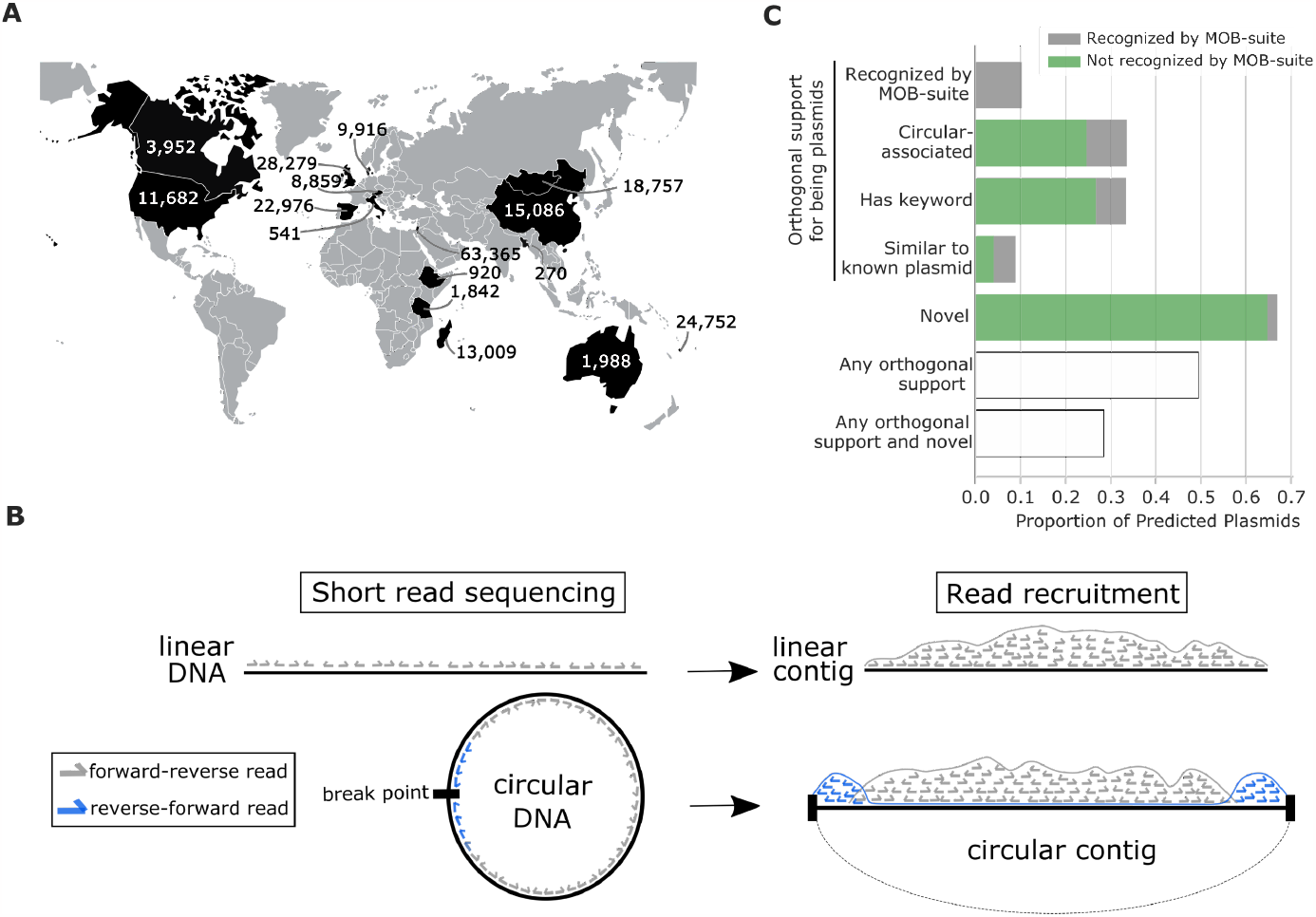
Plasmid prediction from metagenomes. **(A)** Number of plasmids predicted from each country. **(B)** Diagram of paired-end reads mapping to a linear versus a circular contig. Linear contigs have forward-reverse reads only, while circular contigs also have reverse-forward reads concentrated on the ends due to an artifact in contig assembly. **(C)** Orthogonal support for and novelty of the 100,719 non-fragment predictions. We performed several types of analyses to assess how many predictions are true plasmids or novel sequences. While most predictions were not recognized by MOB-suite (gray bars), we found that 24,689 of these (24.5% of the 100,719 non-fragment predictions) were circular-associated sequences. 26,921 (26.7%) were ‘keyword-recognizable’, as they contained a COG or Pfam function with the words ‘plasmid’ or ‘conjugation’. And finally, 3,996 (4.0%) were highly similar to a known plasmid sequence in NCBI, while 65,117 (64.7%) were novel sequences with no hits to any sequence in NCBI. As these different subsets of plasmids are partially overlapping, we took their union to find that 49,739 (49.4%) of predictions had some orthogonal support for being a plasmid, by MOB-suite or any of the first three types of analyses, and 28,658 (28.5%) had such support and were novel.

To determine the circularity of plasmid sequences in metagenomes, we analyzed the orientation of mapped metagenomic paired-end reads (Figure 2B). With this approach we found that 19,652 plasmid sequences were circular, and we designated them as high-confidence plasmids for downstream analyses. Circular plasmids had a median lenth of 4.4 kbp and included sequences that were longer than 25 kbp (n=854), 50 kbp (n=378), or 100 kbp (n=47). An additional 14,151 sequences were not circular themselves but were highly similar to a circular sequence. Together, these two types of sequences defined a set of ‘circular-associated’ sequences representing 33.6%, or 33,803 of 100,719, of the predictions. Multiple factors can explain the lack of signal for circularity for the remaining plasmids, including insufficient sequencing depth to observe a sufficient number of reverse-forward pairs, fragmented contigs, or the non-circular nature of some plasmids that occur linearly^54^ or those that are integrated into chromosomes^3^.

Beyond circularity, confirming *in silico* whether a novel sequence represents a plasmid is a significant challenge. While single-copy core genes have been used to assess the completeness of non-plasmid and non-viral genomes assembled from metagenomes^22^, our understanding of the canonical features of plasmids is limited to a relatively small set of well-studied genes that are primarily derived from plasmids of model organisms in culture^37,38^. For instance, MOB-suite^38^ identified canonical features for plasmid replication and conjugation in only 61% of the reference plasmid subtypes we used to train PlasX, which reveals the limits of conventional approaches to identify plasmid features and survey novel plasmids (Table S1). Indeed, MOB-suite identified canonical features in only 10.1% of our predictions (Table S8). Given this narrow sensitivity, we developed orthogonal data-driven strategies to increase confidence in our predictions (Supplementary Information). We found that 49.4% (49,739) of predictions had orthogonal support for being a plasmid, by MOB-suite or other metrics (Figure 2C), and 28.5% (28,658) had such support and were also novel (Table S8). Overall, these findings suggest that our collection of predicted plasmids include not only sequences that match known plasmids but also novel ones that can further advance our ability to infer the gene pool and ecology of naturally occurring plasmids.

While conducting experiments is the most reliable strategy for validation, the labor-intensive and low-throughput nature of such investigations represent a significant limit to their scale. Nevertheless, to experimentally validate at least some of our metagenome-derived predictions as true plasmids of the human gut, we developed a pipeline to identify predictions that are (1) present in human gut microbial isolates, (2) are circular in those isolates, and (3) can be naturally transferred to other microbes. First, we detected 127 of our predicted plasmids in 14 *Bacteroides* isolate genomes that we sequenced in a previous study^55^ (Figure S4). For two of these plasmids, pFIJ0137_1 and pENG0187_1, we performed additional short-read and long-read sequencing to obtain complete plasmid genomes and confirm their circular configuration (Figure S5). Finally, we demonstrated the ability of pFIJ0137_1 to transfer as a plasmid and confer antibiotic resistance from one *B. fragilis* host to another (see Methods, Figure S4). While not comprehensive, these experimental results show that PlasX is able to predict novel plasmids that have canonical features of being extrachromosomal, circular, and transmissible between cells.

Overall, our survey of individual assemblies of human gut metagenomes using PlasX resulted in 100,719 plasmid sequences for in-depth characterization.

### Novel plasmids are highly prevalent and reflect human biogeography, but their ecology is not explained by microbial taxonomy

Next, we sought to investigate the ecology of plasmids across human populations by creating a non-redundant collection of all plasmids and using metagenomic read recruitment to quantify their distribution across individuals. The dereplication step resulted in 11,121 non-redundant reference plasmids and 68,350 non-redundant predicted plasmids. Recruitment of short reads from the 1,782 globally distributed human gut metagenomes showed that only 1.9% of reference plasmids were present in at least two individuals in our dataset, revealing the limited ecological relevance of reference plasmids to naturally occurring gut microbial communities (Figure 3A). Such a weak detection of reference plasmids in the human gut is likely due to the heavy representation of human-gut associated plasmids in public databases that originate from a relatively small number of human pathogens that are not typically abundant in healthy humans. The reference plasmids did include those that were extremely prevalent across human metagenomes, such as pBI143, a cryptic plasmid that was present in 52% of the gut metagenomes in our dataset, which we investigated in-depth elsewhere (Fogarty et al. 2023). However, the predicted plasmids were much more prevalent across human populations in general (Figure 3B) where 63.1% of them were present in at least two individuals (Figure S2D). In fact, 99.7% of all plasmids that occurred in 100 or more individuals were predicted plasmids.

**Figure 3.**
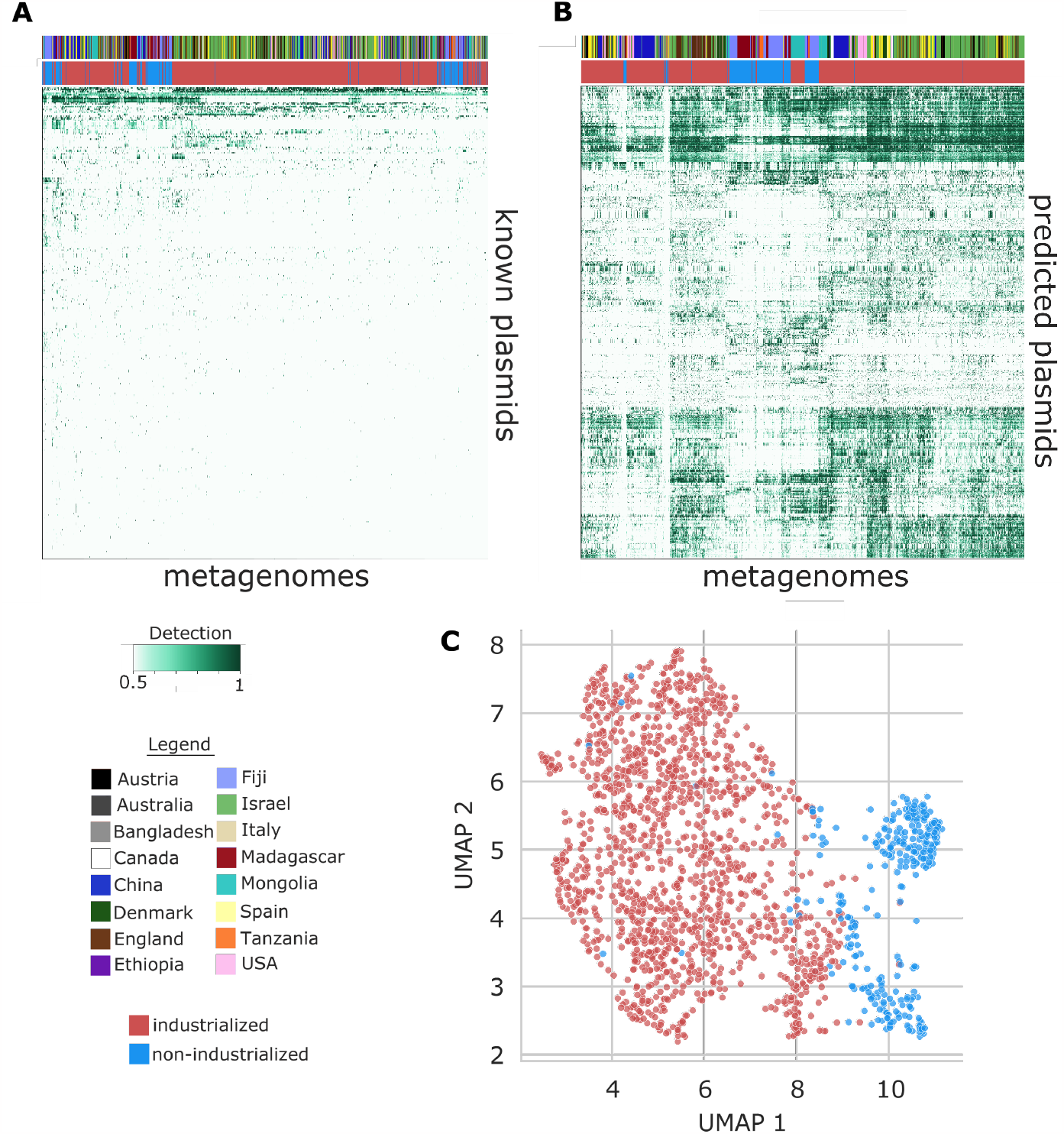
Global plasmid ecology. **(A)** Read recruitment of human gut metagenomes to 11,121 non-redundant reference plasmids. The heatmap shows the 338 plasmids that are present in at least one metagenome (≥0.95 detection). **(B)** Read recruitment to 68,350 non-redundant predicted plasmids. The heatmap shows the 1,000 most prevalent plasmids that are present in at least one metagenome and have PlasX score ≥0.75. In A and B, column colors indicate country of origin and lifestyle (industrialized or non-industrialized). **(C)** Clustering of metagenomes based on the predicted plasmids that are present, using the UMAP dimensionality reduction method^65^. Metagenomes from industrialized or non-industrialized populations are colored red or blue, respectively.

Due to their increased representation in naturally occurring gut microbiomes, plasmids we predicted from metagenomes better capture the biogeography and lifestyles of human populations compared to reference plasmids. Indeed, the organization of metagenomes based on their plasmid content showed that only 50.2% of individuals were placed next to someone from the same country based on reference plasmids (Figure 3A). This percentage increased to 74.0% with plasmids from metagenomes (Figure 3B). Furthermore, their distribution distinguished individuals from industrialized versus non-industrialized countries (Figure 3C), and revealed country-specific clustering of metagenomes (Figure S6).

These results parallel other studies that found associations linking the gut microbial taxonomy with the geography and lifestyles of human populations^56–59^, and thus they lead to an important question: if plasmids and microbial taxa are each correlated with human geography, is microbial taxonomy also correlated with plasmid distribution patterns? On one hand, it would be conceivable to expect a strong correlation between the two as plasmids rely on host machinery for replication and thus their presence in an environment depends on the presence of a suitable microbial taxon. On the other hand, such associations may be weak, or even nonexistent for two reasons: (1) some plasmids are known to have a broad host range^60–62^, and thus their presence in a given environment might not be consistently associated with the presence of a single species or even higher taxonomic category such as a genus or phylum, and (2) plasmids can be gained or lost as a function of environmental pressures, such that nearly identical microbes can differ by the presence or absence of a plasmid or in the number of plasmid copies. By taking advantage of a large number of plasmids that are representative of human biogeography, we examined the ecological associations between plasmid distribution patterns and microbial taxonomy to determine to what extent plasmid ecology can be explained by microbial taxonomy.

For every plasmid, we inferred its most likely host as the taxonomic group that had the most similar ecological distribution (see Methods). While some predicted plasmids had a high ecological similarity with their best matching taxonomic group, the vast majority of predicted plasmids had low similarity scores (median correlation = 0.04, median Jaccard = 0.21) (see Methods, Figures S7A and S7B). We also observed low similarity scores even for reference plasmids that are isolated from a defined microbial host (Figures S7C and S&D). For example, the plasmid pDOJH10S and its cognate host, *Bifidobacterium longum*, were present together in 10 metagenomes; however, in 27 metagenomes we only found the plasmid, and in 69 metagenomes we only found the host (Figure S7E). A more careful detection of these sequences using read recruitment also showed low overlap in metagenomes, with a Jaccard index of 0.40 (Figure S7F, Table S13). The weak correlations between plasmid distribution patterns and microbial taxonomy suggest that plasmids are a highly complex and dynamic feature of microbiomes (Figures 3A, 3B, and S2D), forming an ecological dimension that can stratify human populations (Figures 3C and S6) in ways that cannot be explained by microbial taxonomy alone (Figure S7). While high-throughput analyses of human gut microbiomes often focus on taxonomic features, it has been challenging to find significant or reproducible taxonomic associations that distinguish health and disease states^63,64^. As plasmids often carry key determinants for survival in an environment, a complete understanding of the microbial ecology of health and disease states likely requires the inclusion of insights into plasmid ecology, which requires not only the recovery of naturally occurring plasmids through strategies such as PlasX, but also the characterization of their gene pool in an evolutionary framework, which we aimed to do next using MobMess.

### Plasmid systems organize evolutionarily related plasmids by distinguishing backbone versus cargo content

Our large collection of naturally occurring plasmids provides a unique opportunity to study evolutionary patterns in the human gut plasmidome. Due to frequent genetic rearrangements, a hallmark of plasmid evolution is the reuse of a backbone complemented with variable cargo/accessory genes^13,15,17,18^. Plasmid backbones, which so far have been characterized based on nucleotide identity^13,14,68^, gene similarity^18^, or gene annotations^69–71^, typically encode machinery necessary for plasmid maintenance, while the cargo genes represent additional genetic content, such as antibiotic resistance or other fitness-determining functions.

Here we designed a novel network partitioning algorithm, MobMess (Mobile Element Systems), to study backbone structures in any collection of plasmid sequences at scale (Figure S8). Briefly, MobMess first calculates pairwise alignments across all plasmids to build an initial sequence similarity network, in which a directed edge represents the containment of one plasmid within another, defined by ≥90% sequence identity and ≥90% coverage of the smaller plasmid. We found that these thresholds represent a natural divide between related versus distant plasmids, as we observed a “valley” at these thresholds in the distribution of pairwise average nucleotide identities between all predicted plasmids (see Supplementary Information, Figure S9). Next, MobMess recognizes and collapses redundancy between plasmids, and analyzes patterns of connectivity in the network to identify ‘backbone plasmids’ that satisfy two criteria: (1) the backbone plasmid must be a circular element, inferred here by paired-end orientation (Figure 2B), to ensure that it is not an assembly fragment and can replicate as an independent element, and (2) a backbone plasmid must be found as a subsequence within one or more ‘compound plasmids’. These compound plasmids are composed of the backbone and additional cargo, indicating the ability to acquire or lose genes. Here we define a backbone and its compound plasmids as an evolutionary unit called a ‘plasmid system’ (Figure 4A).

**Figure 4.**
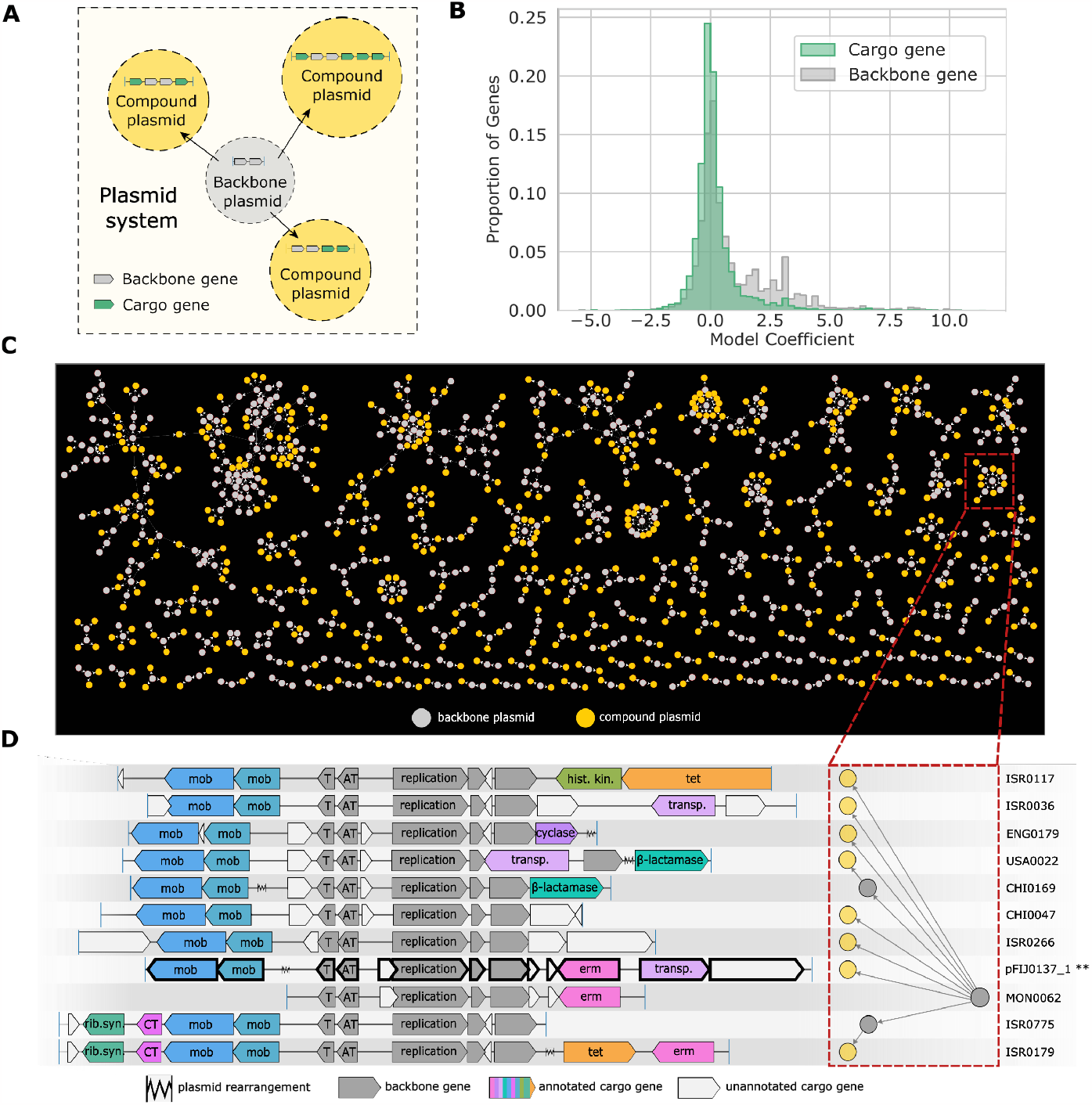
Identification of plasmid systems. **(A)** Network diagram of a plasmid system. **(B)** Distribution of model coefficients for backbone vs. cargo genes in the non-redundant set of 68,350 predicted plasmids. We excluded genes that lacked gene family annotations and thus have a coefficient of zero by default. We also excluded genes that were labeled as backbone with respect to some systems but cargo in others. **(C)** Network of all plasmid systems that contain ≥3 non-redundant and high-confidence plasmids. Only these types of plasmids are shown. **(D)** Genetic architecture of plasmids in PS486, encased by a red box in C. Two plasmids in C are excluded. The system’s backbone (assembled from metagenome MON0062) encodes 5 backbone genes (colored gray). Rib.syn.=riboflavin biosynthesis, CT=conjugative transfer, mob=mobilization, T=toxin, AT=anti-toxin, tet=tetracycline resistance, erm=erythromycin resistance, transp.=transposon, hist. kin.=histidine kinase.

MobMess differs in multiple ways from recently described clustering approaches that have been applied to plasmid sequences^72,73^, which have only been tested on reference plasmids that are complete. In comparison, we designed MobMess to handle complex scenarios by distinguishing between fragmented versus complete (circular) plasmids, a strategy that renders MobMess more suitable to work with complex datasets such as metagenomic assemblies where sequence fragmentation is a common problem. Tracking the containment of smaller plasmids within larger ones is also a significant computational challenge that is overlooked by previous approaches (Supplementary Information). MobMess can distinguish whether a group of plasmids simply share a level of homology through partial alignments or a more nuanced evolutionary relationship through a common backbone that can replicate independently (Figure S10). This way, MobMess minimizes spurious connections between distinct plasmid entities while maximizing its inference of evolutionarily cohesive groups as plasmid systems (Figure S11, Supplementary Information).

This definition of plasmid systems facilitates analyses of plasmid backbone versus cargo content as well as their ecology when used in conjunction with metagenomic data, much in the same way that pangenomes enable studies of core versus accessory gene content in microbial genomes. However, plasmid systems are a specific case of pangenomics, as it is unlikely to find a naturally occurring microbial genome composed only of core genes. In contrast, backbone plasmids represent a minimal entity that can propagate using only backbone genes. With its algorithmic considerations, MobMess provides an automated framework and vocabulary to explore the concept of plasmid systems across different studies and datasets.

### MobMess identifies 1,169 plasmid systems in human gut metagenomes with a wide repertoire of cargo functions

Our application of MobMess to the plasmid sequences predicted by PlasX from human gut metagenomes resulted in a total of 1,169 plasmid systems. Plasmid systems captured a small fraction of the genetic diversity among non-redundant plasmids (6.5%, or 4,424/68,350), however, they captured a large fraction of all circular plasmid contigs (72.7%, or 14,285/19,652) (see Methods, Table S9), and plasmids that were part of a system tended to be longer than plasmids that were *not* part of any system (Table S10). Due to our stringent criteria for the inclusion of plasmids in plasmids systems, MobMess identifies reliable plasmids with multiple representatives in a given dataset, independently of the initial confidence scores assigned by PlasX. For instance, of all plasmids with scores between 0.5 and 0.9, MobMess placed 16,663 in plasmid systems, which retrospectively suggests that a strict cutoff on PlasX prediction scores (such as >0.9) will remove many genuine plasmids from downstream analyses.

Plasmid systems were highly heterogeneous in their genetic complexity. 37 plasmid systems contained sequences that could be classified among 7 different plasmid incompatibility types (Inc11, Inc18, IncFIB, IncFIC, IncI-gamma/K1, IncK2/Z, IncW) (Table S9). 602 plasmid systems contained at least 2 non-redundant compound plasmids, with the largest system containing 168 non-redundant compound plasmids (Figure 4C). For example, pFIJ1037_1, the plasmid we isolated and transferred between *B. fragilis* organisms, was part of PS486, a system containing 24 non-redundant plasmids and found across a total of 127 metagenomes. PS486’s backbone consists of a replication protein and a toxin-antitoxin system, and the cargo genes include beta-lactamases, erythromycin resistance, tetracycline resistance and riboflavin biosynthesis (Figure 4D, Table S11).

To understand how much genetic content is typically conserved or variable in a plasmid system, we calculated the percentage of genes on compound plasmids that were backbone genes versus cargo genes (see Methods). Plasmid systems spanned a wide range of cargo gene percentages between 0% and 100%, with a median value of 40% (Figure S12). Conversely, the median backbone percentage was 60%. PlasX often assigned higher model coefficients to backbone genes in the non-redundant set of predicted plasmids, suggesting these genes define the ‘essence’ of a plasmid by encoding essential functions that promote the ability of a plasmid to exist as a distinct element from the chromosome, such as the genes for plasmid replication, *repA* (PF01051), and mobilization, *mobA* (PF03432) (Figure 4B). In contrast, PlasX assigned lower coefficients to cargo genes, suggesting they encode functions that are not universally essential but important for specific niches, such as nitrogen reductase, *nifH* (PF00142), and membrane transport, *ompA* (PF00691). Indeed, 24.1% (2,169/8,995) of backbone genes versus 13.4% (3,229/24,168) of cargo genes encoded COG and Pfam functions with descriptions related to plasmid replication, transfer, and maintenance (Supplementary Information).

The most frequent type of function encoded on cargo genes was antibiotic resistance, including efflux pumps, which can provide general resistance to multiple antibiotics, and genes targeting specific classes of antibiotics, such as glycopeptides and beta-lactams (Figure 5A). This large-scale observation is consistent with numerous examples of known plasmids encoding resistance and further illustrates how the widespread presence of these plasmids pose a public health threat^76–79^. Other highly prevalent cargo functions included a wide diversity of cellular and metabolic pathways defined in the COG (Figure 5B) and KEGG databases (Figure S13). The most enriched among these was tRNA modification, encoded in 35 compound plasmids within different systems. For example, the globally prevalent system PS1110 (present in 739 metagenomes) contained 291 compound plasmids (27 non-redundant), three of which encoded an enzyme that performs tRNA Gm18 2’-O-methylation (COG0566) and were collectively present in 498 metagenomes (Figure S14, Table S11). This enzyme is thought to reduce the immuno-stimulatory nature of bacterial tRNA, which is detected by Toll-like receptors (TLR7) of the mammalian innate immune system^80,81^. While plasmids in some pathogens are known to facilitate bacterial evasion of mammalian immune system by regulating surface proteins^82^, the overwhelming prevalence of tRNA modification enzymes in our data suggests the likely presence of a previously unappreciated role for plasmids to increase the fitness of their bacterial hosts against the surveillance of the human immune system.

**Figure 5.**
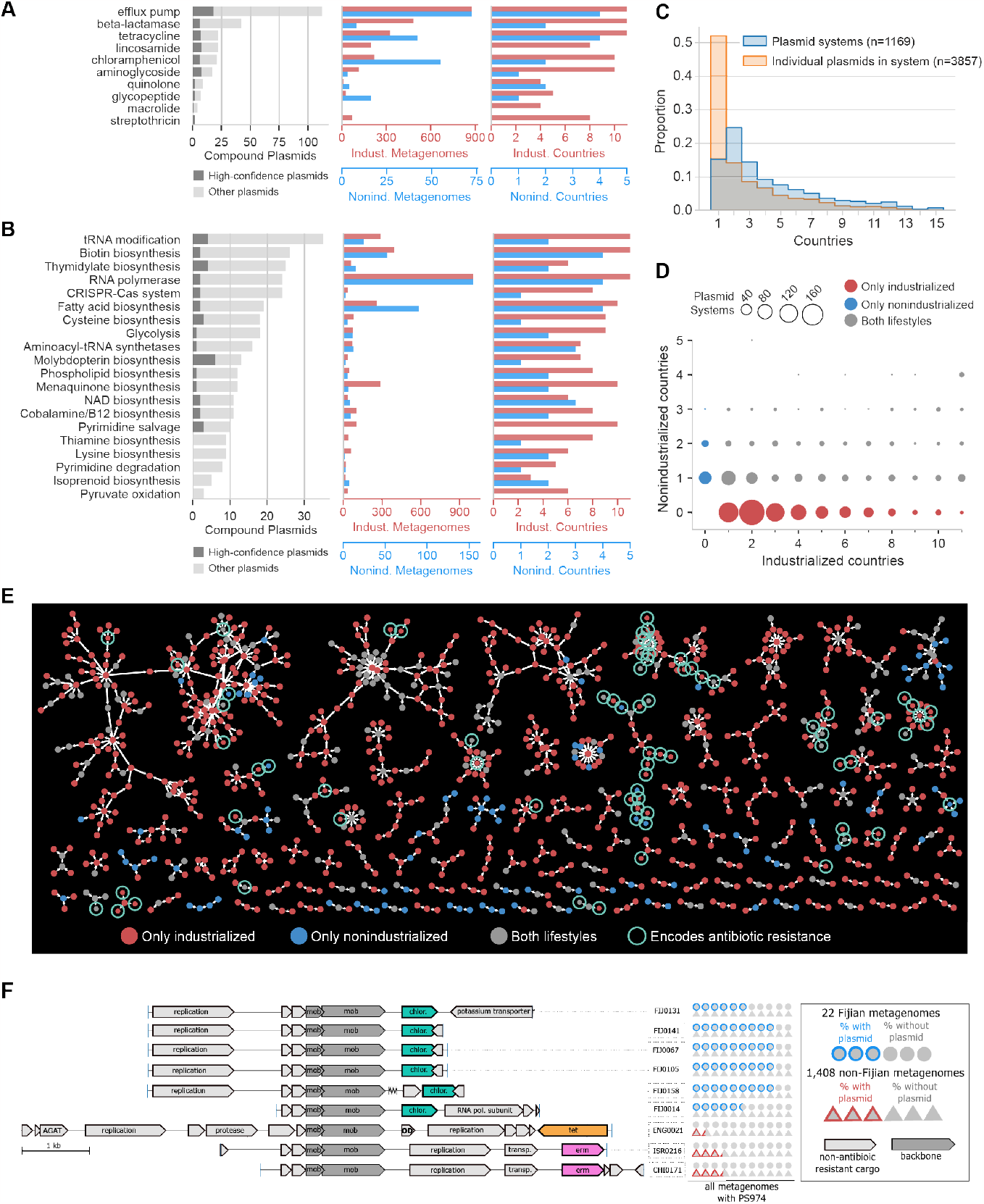
Functional and ecological variation of plasmid systems. **(A-B)** The number of compound plasmids that encode antibiotic resistance (A) and COG pathways (B) in cargo genes. Also shown are the numbers of metagenomes and countries that contain those plasmids. In B, we show the 20 COG pathways that have the highest number of compound plasmids. To avoid redundancy with A, we exclude COG pathways that occur in cargo genes encoding antibiotic resistance. **(C)** Prevalence of plasmid systems versus the individual plasmids in those systems. **(D)** Distribution of plasmid systems based on the number of industrialized and non-industrialized countries they are found in. **(E)** Recoloring of the network of plasmid systems shown in Figure 4C. Colors indicate whether a plasmid occurred in only industrialized, only non-industrialized, or both types of countries. A green ring indicates a plasmid encoding antibiotic resistance. **(F)** Compound plasmids from PS974 that encode for resistance to chloramphenicol (chlor), tetracycline (tet), or erythromycin (erm). 6/9 plasmids are circular. Dark gray genes are the backbone; light gray are cargo not related to antibiotic resistance. AGAT=aminoglycoside adenylyltransferase. OD=Oxaloacetate decarboxylase. PS974 is found in 22 Fijian and 1,408 non-Fijian metagenomes. The pictogram on the right-hand side represents these metagenomes using two shapes: circles (Fijian) and triangles (non-Fijian). For each plasmid, circles are colored blue to represent the proportion of the 22 Fijian-metagenomes that contain the plasmid. Similarly, triangles are colored red to represent the proportion of the 1,408 metagenomes that contain the plasmid.

Overall, these results show that the plasmid systems we were able to identify with MobMess from the human gut represent evolutionarily cohesive units with the enrichment of different classes of functions in the backbone and cargo gene pools. Functional annotations further suggest that while the conserved pool of backbone genes can yield insights into plasmid compatibility and maintenance, the dynamic pool of cargo genes could serve as a means to identify genetic determinants of fitness that respond to particular environmental conditions.

### Plasmid systems reveal cargo genes adapted to specific environments

Given the highly heterogeneous biogeography of individual plasmids (Figure 3B) and their organization into their country of origin (Figure S6), we next asked whether the higher order evolutionary units described by plasmid systems simply consisted of plasmids with similar distribution patterns, or spanned larger geographical regions with individual plasmids of distinct ecology. Our analysis of plasmids and plasmid systems across metagenomes showed that while individual plasmids were often present in a single country, plasmid systems frequently spanned multiple countries (Figure 5C, Table S9). Indeed, of the 2,005 individual plasmids that were unique to a single country, 1,794 (89.5%) were part of more geographically diverse plasmid systems that were present in at least two countries. In fact, 84% (982/1,169) of plasmid systems in our dataset were present in at least two countries, and we found 9 plasmid systems were present in as many as 15 of the 16 countries (Table S9), suggesting that cargo genes that were likely selected for in different regions stemmed from conserved backbone structures.

The broad ecological distribution patterns of plasmid systems compared to the country-specific distribution patterns of plasmids they describe presents a unique opportunity to gain insights into environmental pressures that drive the composition of cargo genes within individual plasmid systems. To investigate this further, we focused on plasmid systems that spanned industrialized and non-industrialized countries, which were largely separated by the distribution of individual plasmids (Figure 3C). Many plasmids systems were exclusive to either industrialized or non-industrialized countries; however, 396 of them were present in both (Figure 5D). Antibiotic usage is a well-understood environmental pressure that often requires microbes to maintain plasmids with antibiotic resistance genes^83–90^. In our data, the evolution of antibiotic resistance in a plasmid system coincided with the ecological variation of compound plasmids in the system. Specifically, we identified 24 high-confidence, compound plasmids that encoded antibiotic resistance in cargo genes and were exclusively present in either non-industrialized or industrialized countries (Figure 5E). Among non-industrialized metagenomes, one of the most common types of antibiotic resistance is chloramphenicol resistance (Figure 5A). For instance, the plasmid system PS974 contained 97 non-redundant plasmids; however, this system possessed chloramphenicol resistance (conferred via an acetyltransferase) only in plasmids from Fiji (Figure 5F, Table S11). When we searched for these resistance plasmids across the global set of 1,430 metagenomes that contain PS974, we found them in 19/22 Fijian metagenomes but only in 1/1,408 non-Fijian metagenomes (*p*=1.1 × 10^−13^, Fisher’s exact test) (Figure 5F, pictogram). Chloramphenicol is routinely prescribed in Fiji to treat eye infections, central nervous system infections, periodontitis, shigellosis, typhoid and paratyphoid fevers, and diabetic foot infections, but it is rarely used in North America and Europe^91–94^. Strikingly, PS974 also contained compound plasmids that carry tetracycline resistance (171/1,408 metagenomes) or erythromycin resistance (429/1,408 metagenomes), yet these plasmids only occurred in individuals from China, Israel, and the United Kingdom (Figure 5F). Matching sequences to these plasmids on the NCBI databases, suggested that their possible microbial hosts include populations in the phylum Firmicutes, such as *Blautia hydrogenotrophica*. However, the distribution patterns of plasmids in PS974 did not match any microbial taxa (highest Jaccard index=0.37 across all plasmid-taxon comparisons). Our observations here suggest that despite sharing a common backbone, compound plasmids in PS974 give access to different antibiotic resistance genes, and their ecology is defined by lifestyle-specific usage of antibiotics.

While the connection between antibiotic usage and resistance is expected given previous studies^83–90^, plasmid systems in general can be used to identify cargo genes that likely determine fitness given known environmental pressures, as well as to hypothesize the existence of environmental pressures given known functions that differentially occur across environments. Overall, our results show that plasmid systems provide a biologically meaningful and computational framework to study plasmid ecology and evolution at scale.

## Discussion

Plasmids are found in nearly every microbial ecosystem, yet the computational challenges associated with their de novo identification have made it difficult for microbiologists to routinely survey plasmids and define evolutionarily cohesive units to describe plasmid diversity in complex environmental samples. PlasX and other plasmid recognition systems^30,31,35–37,45,95–97^, along with MobMess to characterize plasmid systems, present a powerful roadmap for a detailed characterization of naturally occurring plasmids at scale. Even though our study focused on the human gut microbiome, we designed PlasX and MobMess using a broad collection of reference sequences, so that they can be applied to study any environment, and with the flexibility to include additional training sequences to improve accuracy. These methods provide a complementary approach to frequently used state-of-the-art workflows to study the taxonomic composition or functional potential of environmental or host-associated microbiomes through amplicon sequences or metagenomes.

Historically, plasmids and other genetic elements have been characterized on the basis of qualitative properties and descriptions. Early applications of machine learning approaches to predict plasmids employed a small number of marker genes, which limited the recognition of plasmids that lacked recognized markers. The effect of this shortcoming is clear, as many of the genuine plasmids contained no such markers (Figure 2C). Another popular strategy to recognize plasmids has been to employ k-mers, yet such algorithms will also miss plasmids that have low sequence similarity to those that are described in public databases (Figure 1D). By relying upon gene families, PlasX presents an unbiased addition to our bioinformatics toolkit to predict plasmids. But even though both our quantitative and qualitative surveys showed that PlasX surpasses the performance of state-of-the-art algorithms to identify plasmids, it is essential for researchers to take into consideration that predicted sequences will contain both false positives and false negatives. Plasmids can be difficult to distinguish from other mobile or integrated genetic elements as they share common features, including being extrachromosomal^32,98^, facilitating horizontal gene transfer^99,100^, or encoding traditional core functions like replication and mobilization^38,101^. PlasX’s accuracy is also tied to the underlying training set of reference plasmids and chromosomes; over- or under-representation of certain types of sequences can bias the model and limit generalizability.

The confidence scores PlasX assigns to each prediction may serve as a means to adjust the amount of noise in prediction results; however, researchers must consider the trade-off between sensitivity (i.e., capturing a higher fraction of true plasmids) versus specificity (i.e., reducing the number of falsely predicted plasmids) while setting a cut-off. The trade-off between sensitivity and specificity was most visible in our cross-validation analyses, where a threshold of >0.5 lies at an inflection point in the precision-recall curve of Figure 1D (with a precision of 0.850 and recall of 0.500). This threshold was also sufficient at distinguishing ICEs and prophages from plasmids. Applying a stricter threshold of >0.9 increases precision by 13% (to 0.920), but it also decreases recall by 44% (to 0.280). As our understanding of plasmid diversity in metagenomes is greatly underdeveloped, here we decided on applying a threshold of >0.5, in order to provide a reasonable balance between precision and recall for our study and to include many potentially novel plasmids in our results. A good example for this is the novel and long-missed Wolbachia plasmid^47^, which has a score of 0.73. At the same time, stricter thresholds (such as >0.9) or other filters such as circularity may be more appropriate in future work where a higher precision of predicted plasmids (fewer false positives) is a higher priority.

Given the dynamism of plasmids, organizing them into evolutionarily cohesive groups is a significant challenge. Previous computational methods for organizing plasmids have relied on average nucleotide identity (ANI) to represent the whole-sequence similarity between plasmids, but while computational tractable, this single statistic misses other evolutionary dimensions that relate sequences. To address these issues we developed MobMess, which resolves the containment of plasmid sequences within one another. This methodological advance enabled us to identify plasmids systems, revealing the great extent to which plasmids in complex ecosystems are not static entities but actively evolving in response to the environment. Plasmid malleability is a desirable property in bioengineering and has often motivated the repurposing of naturally occurring plasmids into major tools for genetically modifying organisms. In this vein, we propose computational identification of plasmid systems as an attractive approach to expand the toolkit of available plasmids for genetic engineering, particularly if they are found in isolates that currently lack tools to make them genetically tractable. Plasmid systems, which manifest in many distinct forms across multiple human populations, excel at incorporating new functions and propagating across a wide range of natural environments and may behave similarly in laboratory settings.

An overarching implication of our findings is that high-throughput recognition and characterization of plasmids in microbiome studies are necessary for more complete insights into the ecology of naturally occurring microbial systems.

## Methods

### Compiling and annotating a reference set of plasmids and chromosomes

We obtained a list of 16,168 plasmids from the 2019_03_05 version of PLSDB^102^. We also downloaded the entire collection of 13,471 complete bacterial genome assemblies from NCBI RefSeq on October 26, 2019, using instructions at https://www.ncbi.nlm.nih.gov/genome/doc/ftpfaq/#allcomplete. The RefSeq assemblies contained 26,376 contigs, of which we discarded 11,350 that are also in PLSDB. The reference set of 16,827 plasmids consisted of 16,168 PLSDB contigs, as well as 659 contigs from the Refseq assemblies that were labeled as ‘Plasmid’ in the ‘Assigned-Molecule-Location/Type’ field of the NCBI assembly report. The reference set of chromosomes was the remaining 14,367 Refeq contigs.

To identify and annotate genes in these sequences, we used the program ‘anvi-run-workflow’ with ‘--workflow contigs’ implemented^103^ in anvi’o^104^ v7.1, which uses Snakemake^105^ to execute previously defined steps (https://merenlab.org/anvio-workflows/) to generate anvi’o contigs-db files (https://anvio.org/m/contigs-db). These steps include first running Prodigal^106^ to call genes and then running DIAMOND v2.0^107^ and HMMER v3.3^108^ on amino acid sequences to determine gene functions against the Cluster of Orthologous Groups of proteins (COGs)^42^ and Protein Family Database models (Pfams) v32.0^43^, respectively. To minimize noise, we used an e-value cutoff of 10^−10^ for COGs and the default model scores for Pfams.

### Modeling *de novo* gene families

We inferred *de novo* gene families by running MMseqs2^109^ v10.6d92c on all amino acid sequences in our reference plasmids and chromosomes. First, we ran ‘mmseqs clusthash’ to collapse identical sequences into a non-redundant set for faster execution of the next step; the collapsing was inverted at the end to annotate all genes. Next, we ran ‘mmseqs cluster’ to calculate pairwise alignments and then cluster genes that are aligned above a minimum sequence identity threshold (parameter ‘--min-seq-id’). We ran this program multiple times with different thresholds (0.9, 0.8, 0.7, 0.6, 0.5, 0.4, 0.3, 0.25, 0.2, 0.15, 0.1, 0.05) to infer a wide range of possible families. Families from different thresholds can be redundant, so we merged nested families, i.e. if family X contains all genes in family Y, then we keep X and discard Y. We also discarded any family that contains only one gene. In theory, families inferred from a higher threshold (e.g. 0.9) should always nest within a family inferred from a lower threshold (e.g. 0.05), such that we would discard all families from higher thresholds. But in practice, families don’t always nest within each other but only overlap partially. After merging, our final model used the following number of families from each threshold.

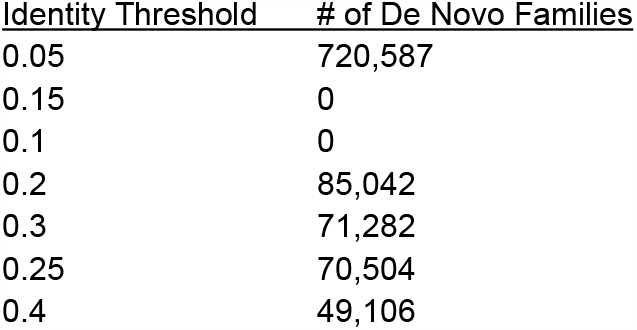
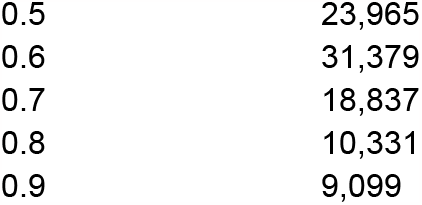

In total, our model used 1,090,132 gene families, which annotated 162,783,114 genes. Note that because these gene families can still overlap with each other, a gene may have multiple annotations. This analysis took advantage of MMseqs2’s parallelism, taking ∼6 hours using 256 CPU cores.

We refer to a *de novo* family as a ‘subfamily’ if 90% or more of its set of amino acid sequences are also annotated to a specific COG or Pfam. Note that this definition provides a small tolerance such that a subfamily does not need to be a perfect subset of a COG or Pfam. For the example about Pfam PF10609 (Figures 1F and 1G), we gathered the 253 amino acid sequences annotated to PF10609 and the subfamily mmseqs_5_1535552. We also gathered the 1,391 sequences annotated to PF10609 and the subfamily mmseqs_70_40217271. We collapsed 100% identical sequences to yield a total collection of 142 and 310 sequences from mmseqs_5_1535552 and mmseqs_70_40217271, respectively. We aligned all of these sequences together using muscle v3.8.1551 (default parameters)^110^ and then constructed a maximum likelihood phylogenetic tree using IQ-TREE v2.1.2 (parameters -m TEST -bb 1000 -alrt 1000 -T AUTO)^111^. We then rooted the tree using the midpoint method.

### Subtypes and slicing of reference sequences

To group reference sequences into subtypes, we used mash v2.2.2^112^ (command ‘mash dist’, sketch size 100000, kmer size 21) to calculate a distance score of 0 to 1 between every pair of sequences. Next, we created an undirected graph, where sequences are nodes and sequences are connected if their distance is ≤0.1. We defined a ‘subtype’ as one of the 7,326 connected components in the graph. 3,935 subtypes contained only plasmids; 3,355 subtypes contained only chromosomes; and 36 subtypes contained both plasmids and chromosomes (Supplementary Table S1).

We sliced reference sequences into 10 kbp slices by sliding a 10 kbp window at 5kb increments. The first window starts at the beginning of the sequence, and the final window stops at the end of the sequence. For instance, a 23kb sequence would be sliced at 0-10 kbp, 5-15 kbp, 10-20 kbp, and 13-23 kbp. A slice was annotated with any gene that was entirely or partly inside the slice. In total, we generated 10,453,279 slices from the reference chromosomes and 343,246 slices from the reference plasmids.

### Assessing model performance in cross-validation

We performed 4-fold cross validation by splitting the set of 10 kbp slices of reference sequences into four random subgroups. We used the sequences in three subgroups to train our model, PlasX, and then we evaluated the performance of the model on the fourth subgroup. We redo this procedure by changing which subgroups are used for training or for evaluation – for a total of four times. This procedure is a common technique in machine learning, more generally known as “k-fold validation” where k is the number of subgroupings. In a naive split, we keep all slices from the same reference sequence together in either training or testing data. In an informed split, we keep all slices from the same subtype together.

We assigned weights to the 10 kbp slices when calculating precision and recall performance (Figure S1F and 1D). Consider the following notation to represent sequences:

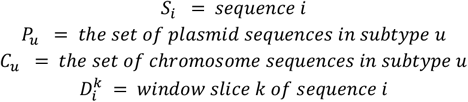

And consider the following notation to represent weights:

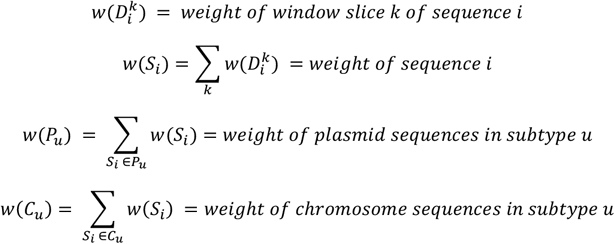

We defined two different scenarios for assigning weights. Scenario A satisfies the following conditions:

1. All slices from the same sequence have the same weight

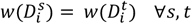
2. The weight of every sequence is equal to 1

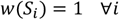

Scenario B satisfies the following conditions:

1. All slices from the same sequence have the same weight

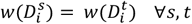
2. All plasmid (or chromosome) sequences in the same subtype have equal weight

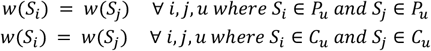
3. All subtypes have equal weight

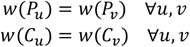
4. The sum of weights across all slices equals the total number of slices

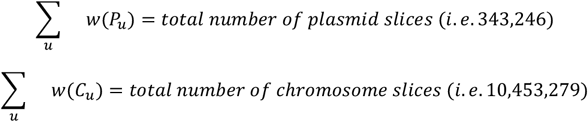

Each scenario implies a unique assignment of weight values. Scenario A requires that every sequence has the same weight. Importantly, this ensures that long sequences, which have disproportionately more slices, have equal weight as shorter sequences. Scenario B further requires that every subtype has the same weight. Importantly, this ensures that subtypes that contain a disproportionately large number of sequences (e.g. subtypes that represent commonly studied bacteria, such as *Escherichia, Salmonella*, and *Klebsiella*) have equal weight as subtypes with fewer sequences.

We evaluated performance under two different cross-validation and weighting scenarios. Figure S1F shows the result of training models using a ‘naive’ cross-validation split and calculating precision/recall using weights from Scenario A. Figure 1D shows the results of training models using an ‘informed’ cross-validation split and calculating precision/recall using weights from Scenario B. We calculated precision/recall using the function *sklearn*.*metrics*.*precision_recall_curve* from the scikit-learn Python package^113^, with the parameter *sample_weight* set to the weights of the slices. We calculated AUCPR with the function *sklearn*.*metrics*.*average_precision_score*.

### PlasX implementation

We implemented PlasX as a logistic regression using the *SGDClassifier* class from scikit-learn^113^. For training and evaluating PlasX, we used 10 kbp slices of the reference sequences, to normalize for the fact that chromosomes are generally much longer than plasmids and to improve downstream application of PlasX on sequence collections that may contain a large number of fragmented sequences, such as assembled metagenomes. Regardless of how we evaluated PlasX (either Figure S1F or Figure 1D), we always trained it with weights defined by Scenario B and based on only slices in the training data.

To improve performance, PlasX uses a technique called elastic net regularization, which identifies gene families with redundant or noisy signals and then minimizes the usage of these families by setting their coefficients equal or close to zero. Consequently, only a non-redundant and informative set of gene families can impact predictions by having coefficients far from zero (Figure S1D). To implement elastic net regularization, we performed a grid search of hyperparameters, with the regularization parameter *alpha* ranging from 10^−8^ to 10^−3^ in multiplicative increments of 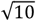 and the parameter *l1_ratio* being 0.0, 0.25, 0.5, 0.75, or 1.0. For each evaluation scenario (either Figure S1F or Figure 1D), we selected the hyperparameters that produced the best performance. We used the best hyperparameters from the ‘informed’ cross-validation and the weights defined by Scenario B (*alpha*=3.16x10^−6^, *l1_ratio*=0.0) to retrain PlasX on all 10 kbp slices and create the final model that we used to predict plasmids from metagenomes.

### Predicting plasmids from metagenomic assemblies

We downloaded fastq files for 1,782 short-read and paired-end metagenomes from the National Center for Biotechnology Information (NCBI) Sequence Read Archive (SRA) using the program ‘fastq-dump’. The countries represented are Austria^114^, Australia (https://www.ncbi.nlm.nih.gov/bioproject/PRJEB6092), Bangladesh^115^, Canada^116^, China^117,118^, Denmark^119^, England^120^, Ethiopia^58^, Fiji^121^, Israel^122^, Italy^123^, Madagascar^58^, Mongolia^124^, Spain^125^, Tanzania^123^ and USA^126,127^. Some samples were sequenced multiple times (i.e. multiple records in SRA), in which case we concatenated the fastq files together. We have separated these multiple accessions using the delimiter ‘|’. We labeled Tanzania, Ethiopia, Bangladesh, Madagascar, and Fiji as non-industrialized and the other countries as industrialized.

All steps of quality filtering, metagenomic assembly, read recruitment and profiling were automated using snakemake^105^ workflows in anvi’o^128^. The ‘illumina-utils’^129^ commands ‘iu-gen-configs’ and ‘iu-filter-quality-minoche’ with the flag ‘--ignore-deflines’ were used to quality filter the raw paired-end reads. Each metagenome was assembled individually using IDBA_UD^130^ with default settings, except the flag ‘--min_contig 1000’.

We annotated COGs and Pfams in all assembled contigs using the same procedure as the reference plasmids and chromosomes. To annotate *de novo* families, we first used ‘mmseqs result2profile’ (default parameters) to represent the sequence conservation in each *de novo* family as a profile. We then used ‘mmseqs search’ (default parameters) to search for profiles across all genes. We kept hits where the alignment coverage was ≥80% of both the gene and the profile and where the alignment identity was at least ≥X-0.05 where X is the minimum identity threshold used to originally construct the family (parameter --min-seq-id). For example, if a family was constructed using an identity threshold of 0.8, then we kept hits with an identity ≥0.75. Using these gene annotations, we ran PlasX to assign a score to every contig. We kept contigs intact, rather than slicing them into 10 kbp windows. Contigs with score >0.5 were classified as plasmids.

### Detection and circularity of plasmids across metagenomes

We recruited short reads from our collection of metagenomes using Bowtie2 v.2.0.5^131^. We used the snakemake workflows in anvi’o to automate execution of bowtie and post-processing to calculate ‘detection’, i.e. the proportion of a sequence that is covered by at least one read. We ran bowtie2 using the three following combinations of parameters and input files.

First, to identify circular contigs, we recruited each metagenome’s reads to a fasta file that contained only the contigs assembled from that metagenome. For computational efficiency, we ran bowtie2 with its default behavior to align every read at most once. We then analyzed the orientation of paired-end reads (Figure 2B). During assembly, circular sequences are broken by an artificial breakpoint to represent them as linear contigs. Consequently, DNA sequencing that occurred across this breakpoint will produce paired-end reads that align in a reverse-forward orientation to the ends of the contig. In contrast, if a sequence is not circular, then all paired-end reads are expected to align in a forward-reverse orientation. To illustrate this intuition, suppose the upstream read of a paired-end maps to positions 200-300 of a contig and the downstream read maps to 500-600. If the upstream read maps with a reverse complement strandedness (i.e. ‘reverse’) and the downstream read maps with the same strandedness as the way the contig is written (i.e. ‘forward’), then the paired-end is in a reverse-forward orientation. In other words, if the contig is written 5’-to-3’, then the upstream read maps 3’-to-5’ and the downstream read maps 5’-to-3’. Inversely, the paired-end is in a forward-reverse orientation if the upstream read maps 5’-to-3’ and the downstream read maps 3’-to-5’. Next, we defined the gap (or insert) size of a paired-end to be the distance between the closest (or farthest) aligned positions between its two reads. In our example, the gap size is 600-200=400 and the insert size is 500-300=200. Let *D* be the contig’s length minus three times the median insert size of all forward-reverse paired-ends that are aligned to the contig. Finally, we label a contig as circular if (1) its detection was ≥0.95 and (2) it had at least one reverse-forward paired-end with a gap greater than or equal to *D*. This approach of examining reverse-forward paired-ends was inspired by ^132^. There were 154,680 contigs that were not predicted to be plasmids but still appeared circular; however, these contigs tended to have a smaller number of supporting reverse-forward reads relative to their coverage (Figure S2E), which may indicate that they are other types of mobile elements such as viruses or ICEs that temporarily circularize.

Second, to study the ecological distribution of all plasmids and plasmid systems at the same time, we recruited each metagenome’s reads to a fasta file that contained either the non-redundant set of 68,350 predicted plasmids or the non-redundant set of 11,121 reference plasmids. For computational efficiency, every read was aligned at most once (i.e. the default behavior of bowtie). We designated a plasmid as being present in a metagenome if its detection was ≥0.95. To compare metagenomes based on their plasmid content in Figure 3 and Figure S6, we ran UMAP v0.5.1^65^ with parameters ‘n_neighbors=30, n_components=2, min_dist=0.15, metric=‘jaccard’, random_state=1’. The heatmaps in Figure 3 were generated using the ‘heatmap.2’ package in R, with agglomerative clustering using median linkage on Euclidean distances.

Third, to study the specific plasmids from PS974 and PS1110 that were shown in Figure 5F and Figure S14 (see Table S11 for contig names), we ran bowtie2 on each sequence separately. This setup allowed every read to align to potentially multiple sequences, resulting in a more complete estimation of which metagenomes contained a plasmid. For the backbone sequences of these systems, we designated them as present in a metagenome if their detection was ≥0.95. For compound plasmids in PS1110 that encoded a chloramphenicol resistance gene, we designated them as present in a metagenome if they satisfied an additional criterion that ≥0.95 of the resistance gene was covered by at least one read.

### Estimation of potential hosts for plasmids in metagenomes

We used two different formulas to calculate the ecological similarity between a plasmid and potential host: (1) the Pearson correlation in the abundance levels of the plasmid and host across metagenomes, and (2) a Jaccard index to represent the fraction of metagenomes that contain both the plasmid and host.

We estimated taxonomic abundances in every metagenome by running kraken2^133^ v2.1.2 with its standard database (https://github.com/DerrickWood/kraken2) and then refined the abundances using bracken^67^ v2.5 (https://github.com/jenniferlu717/Bracken) with database parameters ‘-k 35 -l 96’. We ran bracken (parameter ‘-r 96’) a separate time for every taxonomic rank: S1 (subspecies/strain), S (species), G (genus), F (family), O (order), C (class), P (phylum), D (domain). The output of this analysis is a count of how many reads originated from each taxon.

To compare the metagenomic presence/absence of plasmids versus taxa, we calculated *M*_*P*_, the set of metagenomes where a plasmid *P* is detected at ≥95% (based on recruitment to the 68,350 non-redundant plasmids), and 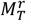, the set of metagenomes in which at least *r* reads originated from taxon *T* (based on kraken and bracken). For each plasmid, we attempted to find the best explanation of its ecological distribution by comparing the plasmid to every taxon using the Jaccard index, and by scanning many possible read thresholds. More exactly, we used the following formula to represent the best possible explanation of a plasmid’s ecological distribution:

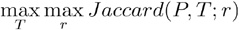

where

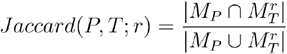

We evaluated 29 values for the threshold *r*, ranging from 1 read to 10 million reads in multiplicative increments of 10^1/4^. We ignored plasmids that were present in less than 5 metagenomes, i.e. |*M*_*P*_|<5, because it was likely that these plasmids would have a high Jaccard similarity to some taxon by random chance. For instance, we observed that many pairs of plasmids and taxa occur in exactly one and the same metagenome, and thus they have a Jaccard index of 1.

To compare continuous-valued abundances, we defined a plasmid’s abundance in a metagenome as the sum of coverage values across all sequence positions divided by sequence length, and we defined a taxon’s abundance as bracken’s estimate of the number of reads originating from the taxon. If a plasmid had detection of <95%, then we set its abundance to 0. If a taxon had less than 1000 reads, then we set its abundance to 0. We ignored plasmids and taxa that had non-zero abundances in less than 5 metagenomes. For every pair of plasmid and taxon, we estimated the Pearson correlation between their abundance levels across metagenomes using FastSpar^134^ v1.0.0 (https://github.com/scwatts/fastspar), which is an improved implementation of the SparCC^66^. This method accounts for the compositional nature of the data— in which abundances reflect relative instead of absolute quantities—by assuming that the amount of correlations in a data is sparse. We ran FastSpar on the non-redundant set of predicted plasmids, and ran it separately on the non-redundant set of reference plasmids.

We performed a more careful analysis of plasmid pDOJH10S and its cognate host *B. longum DJO10A* by performing a read recruitment of metagenomes to both sequences together, using bowtie2 and allowing every read to align at most once (Supplementary Figure S7F, Table S13). Following Utter et al.^135^, we applied a >50% detection threshold to identify the presence of the plasmid or host. 365 metagenomes contained the plasmid, 818 metagenomes contained the cognate host, and 336 metagenomes contained both genomes (Jaccard index = 0.40).

### Keyword analysis of COGs and Pfams for plasmid functions

We labeled COGs and Pfams as being a plasmid-associated function (Figure 1E) if its database description contains any of following keywords as a substring: *‘plasmid’, ‘toxin’, ‘replicat’, ‘integrase’, ‘transpos’, ‘recombinase’, ‘resolvase’, ‘relaxase’, ‘recombination’, ‘partitioning’, ‘mobilis’, ‘mobiliz’, ‘type iv’, ‘conjugal’, ‘conjugat’, ‘segregat’, ‘MobA’, ‘ParA’, ‘ParB’, ‘BcsQ’*. We labeled backbone and cargo genes as being related to plasmid replication, transfer, or maintenance if they were annotated to any plasmid-associated COG or Pfam (see section “Classification of cargo and backbone genes”).

To determine if a predicted plasmid is ‘keyword-recognizable’ (Figure 2C), we searched the plasmids for COGs and Pfams using a more restricted set of keywords (just “plasmid” and “conjugation”) instead of the keywords above.

### MobMess algorithm to dereplicate plasmids, remove assembly fragments, and discover plasmid systems

The MobMess algorithm performs three tasks. It de-replicates plasmids that are nearly redundant to each other; it removes plasmids that appear to be assembly fragments; and finally it organizes plasmids together into evolutionary groups called plasmid systems. MobMess consists of several steps described below.

MobMess first performs an all-vs-all pairwise alignment of sequences using the MUMmer alignment package (v4.0.0rc1)^136^. All sequences are placed into a single fasta file and then aligned with ‘nucmer’ (parameters ‘--maxmatch --minmatch=16’) to calculate local alignment blocks. Alignments are specified asymmetrically such that one sequence is designated as the query *q* and the other is the reference *r*. For every *q* and *r*, the alignment blocks calculated by ‘nucmer’ are written to a separate file, and then a subset of blocks is identified using ‘delta-filter’ (parameters ‘-q -r’) to create a one-to-one alignment.

Next, MobMess constructs a directed graph *G* where vertices are sequences and edges represent the containment of one sequence within another (Figure S8A). Formally, consider a query *q* and reference *r*. Let |*q*| be the length of *q*. For the *i*^th^ alignment block between *q* and *r*, let *s*^*i*^, *e*^*i*^, and *δ*^*i*^ be the start position in *q*, end position in *q*, and number of alignment mismatches and indels, respectively. The following values summarize the information across all alignment blocks between *q* and *r*.

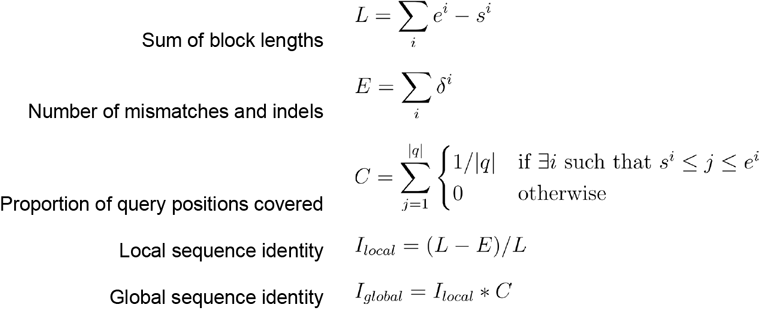

MobMess creates a directed edge (*q,r*) in *G* if *I*_*local*_ and *C* are above user-specified thresholds. In this study, we applied thresholds of *I*_*local*_≥0.9 and *C*≥0.9 (see Supplementary Information). In Figure S9, we re-ran MobMess using various thresholds on *I*_*local*_ and *C* (the same threshold was applied to *I*_*local*_ and *C* at the same time).

MobMess clusters sequences according to strongly connected components in *G*, calculated with igraph v0.8.2^137^ in Python. That is, two sequences *x* and *y* are placed in the same cluster if there exists a directed path from *x* to *y* and another from *y* to *x* in *G*. Intuitively, a cluster represents a set of sequences that are nearly identical to each other across nearly their entire lengths. MobMess then reduces *G* to another graph *H*, called the condensation graph, by contracting every cluster of sequences into a single vertex. A directed edge (*u,v*) exists in *H* if and only if there are sequences *x* ∈ *u* and *y* ∈ *v* where edge *(x,y)* exists in *G*. Note that *H* does not have any cycles. As proof by contradiction, if there were a cycle of clusters, then those clusters would have been in the same strongly connected component in *G* and hence would have been merged into a single, larger cluster.

MobMess labels every cluster in *H* as one of the three following types: (1) a ‘backbone cluster’ if it has an outgoing edge and at least one of its member sequences is circular, (2) a ‘fragment cluster’ if it has an outgoing edge but none of its member sequences are circular, or (3) a ‘maximal cluster’ if it does not have any outgoing edges. Intuitively, a maximal cluster represents the longest version of a plasmid observed in the data. In contrast, a backbone or fragment cluster represents a set of plasmids that are subsequences of other plasmids in a maximal cluster. The only difference between backbone and fragment clusters is that backbone clusters contain at least one circular plasmid (implying complete assembly), while fragment clusters do not contain any circular plasmids (suggesting they are assembly fragments of the maximal cluster).

To dereplicate sequences, MobMess discards all fragment clusters and then chooses a representative sequence from every maximal and backbone cluster. A cluster’s representative is the sequence with the highest global sequence identity (*I*_*global*_), averaged across the set of alignments where that sequence is the reference and other sequences in the same cluster are the queries.

MobMess defines a plasmid system as a specific backbone cluster together with its ‘compound’ clusters, which are the set of non-fragment clusters connected to the backbone in *H*. Thus, there is a one-to-one correspondence between backbone clusters and plasmid systems. Note that systems can be nested within each other, because backbone clusters can be connected to each other in *H*. Thus, a backbone cluster can be *the backbone* that forms a given plasmid system, and at the same time, it can also be a *compound cluster* with respect to an even smaller backbone that forms a different system. As another note, a maximal cluster can be a ‘compound’ cluster of a system, but it is also possible that some maximal clusters are not found in any system because they are not connected to any backbone clusters in *H*.

We ran MobMess to analyze the 226,194 predicted plasmid contigs. MobMess grouped the contigs into a total of 132,616 clusters. 64,266 clusters were ‘fragment clusters’ that contained 125,475 contigs, which we interpreted as assembly fragments of other predicted plasmids. We discarded these fragments from further analysis. The other 68,350 clusters were non-fragment clusters (i.e. 1,169 backbone and 67,181 maximal clusters) and contained 100,719 contigs. Finally, MobMess identified 1,169 plasmid systems, which together represent 1,169 backbone and 63,926 maximal clusters (3,255 maximal clusters were excluded). See Figure S3 for a diagram of these numbers.

We ran MobMess separately on the 16,827 reference plasmid sequences, yielding 11,121 clusters. We assumed that all reference plasmids were circular, and thus there were no fragment clusters. We visualized networks with Cytoscape^74^ v3.8 and laid nodes out using the prefuse directed force layout^75^. While we have focused on plasmids, MobMess could be applied to dereplicate and organize other mobile genetic elements into systems.

### Classification of cargo and backbone genes

We classified all genes on the backbone plasmids of a plasmid system as backbone genes. For genes on compound plasmids, we tested whether the genes shared any *de novo* family annotations with the genes on the backbone plasmids. If so, we classified those genes as backbone genes, otherwise as cargo genes. For this analysis, we used the 1,090,132 *de novo* families that we constructed from reference plasmids and chromosomes in order to train PlasX, and we also used an additional set of 439,584 *de novo* families that we constructed by running the command MMseqs2 (--min-seq-id 0.05) on the genes from all plasmid sequences in this study (16,827 reference and 226,194 predicted plasmids). These additional families allowed us to capture gene families that might be absent in reference sequences but are conserved in predicted plasmids. Note that the classification of genes as backbone or cargo depends on which plasmid system is being considered. It is possible for a gene to be classified as a backbone gene with respect to one plasmid system and, at the same time, as a cargo gene with respect to another system. This is because a plasmid can be a backbone plasmid of a system and also be a compound plasmid of a different system.

For every non-redundant compound plasmid in the system, we calculated the fraction of genes in the plasmid that were cargo genes. We then averaged this fraction across all non-redundant compound plasmids in the system to define the “cargo gene percentage” of the system (Figure S12). Because every gene is either backbone or cargo, the percentage of backbone genes is 100% minus the cargo gene percentage.

For Figure 4B and to analyze the content of backbone/cargo genes, we used a non-redundant and unambiguous set of 8,995 backbone and 24,168 cargo genes. To derive these sets of genes, we first considered the 47,172 genes encoded on the 4,424 non-redundant plasmids that were part of at least one plasmid system. Of these 47,172 genes, we used the 8,995 genes that were classified as backbone genes because they were encoded on a backbone plasmid and that were never classified as cargo genes in any plasmid system. 24.1% (2,169/8,995) of these genes had a plasmid-associated keyword in their COG/Pfam annotations (see Methods section “Keyword analysis of COGs and Pfams for plasmid functions”). We also used the 24,168 genes that were always classified as cargo genes and never backbone genes in any plasmid system. 13.4% (3,229/24,168) of these genes had a plasmid-associated keyword. We excluded from analysis the 1,917 genes that were sometimes classified as backbone genes and other times cargo genes, depending on the system. We also excluded 12,092 genes that were on compound plasmids but were classified as backbone genes, as these genes are redundant with the backbone genes that were encoded on the backbone plasmid.

### Identification of antibiotic resistance genes

We annotated antibiotic resistance genes using two databases. First, we searched against a database of resistance protein family HMMs from Resfams^138^ (v1.2, dated 2015-01-27, ‘Core’ database at http://www.dantaslab.org/resfams). We used ‘anvi-run-hmms’ from anvi’o^104^ to automate running ‘hmmsearch’ from HMMER^108^ 3.3.2 and apply an e-value cutoff of 10^−10^. Second, we ran rgi (v5.2.0, https://github.com/arpcard/rgi) to search for similarity in the CARD database of resistance genes^139^. We removed CARD hits that were labeled as ‘Loose’ and kept those labeled as ‘Perfect’ or ‘Strict’. We removed any Resfams or CARD hits that contained the keywords ‘transcription’, ‘regulat’, ‘modulat’ in their database description, to avoid cases (e.g. TetR protein) where the hit is a gene that regulates the expression of another resistance gene but doesn’t itself perform the molecular process that confers resistance. We categorized hits into major antibiotic resistance classes by searching for the following keywords in their functional descriptions: ‘lincosamide’, ‘macrolide’, ‘erythromycin’, ‘chloramphenicol’, ‘aminoglycoside’, ‘streptothricin’, ‘glycopeptide’, ‘efflux pump’, ‘beta-lactamase’, ‘nitroimidazole’, ‘tetraycyline’, ‘quinolone’, ‘sulfonamide’. Additionally, we searched the extra keywords ‘Van’ and ‘VanZ’ to identify glycopeptide resistance; ‘efflux’, ‘permease’, and ‘pump’ to identify efflux pumps; and ‘TetX’ to identify tetracycline resistance.

### High molecular weight (HMW) DNA extraction, long-read sequencing, and determination of circularity through long-reads

We employed a long-read sequencing strategy on two *Bacteroides fragilis* cultivars from two patients (p-214 and n-216 previously described in Vineis et al^55^). We extracted total genomic HMW DNA by one of two methods. For *B. fragilis* p-214, we used the Qiagen Genomic Tip 20/G procedure (also known as Method #4/GT) as previously described^140^ on a 10 mL overnight BHIS broth culture. For *B. fragilis* n-216, we used a Phenol Chloroform protocol on 25 mL overnight BHIS broth cultures. Libraries were prepared with the Rapid Barcoding Kit (SQK-RBK004) and the standard protocols from Oxford Nanopore Technologies (ONT) with a few modifications. For *B. fragilis* p-214, DNA fragmentation was performed on 6 ug DNA using 5 passes through a 22G needle in a 30 μL volume. The gDNA input was 1.5 μg (Table S12), based on sample availability in a 7.5 μL volume, with 2.5 μL Fragmentation mix added. We sequenced for 72 hours using a single R9.4/FLO-MIN106 flow cell (ONT). For *B. fragilis* n-216, DNA fragmentation was performed on 10 ug DNA using 10 passes through a 22G needle in a 250 μL volume. The gDNA input was 0.32 to 0.44 μg, based on sample availability in an 8.5 μL volume, with 1.5 μL Fragmentation mix added per sample. We sequenced for 72 hours using a single R9.4/FLO-MIN106 flow cell. We used Guppy (v4.0.15) for all post-run base calling, sample de-multiplexing and the conversion of raw FAST5 to FASTQ files.

To determine circularity, we used BLAST to align the long reads with a minimum quality score of 7 to our predicted plasmid sequences following a previously described approach^47^. During assembly, all DNA short reads are assembled as linear sequences even if they are circular elements. Circular elements have an artificial breakpoint to represent them as linear sequences, and this breakpoint can happen anywhere on the sequence depending on the assembly method. We identified and manually confirmed 500 long reads that align completely to a plasmid but not to the host chromosome (Figures S4A and S5B). We tested for the presence of an artificially introduced breakpoint by visualizing these alignments on the sequence as if it were assumed to be a circular element (Figure S5A). If indeed the sequence is circular, the long reads would overlap each other and “wrap around” the entire circumference of the sequence. In other words, all nucleotide positions of the sequence would be covered by at least one read and there would also exist a read that spans the breakpoint by aligning to both sides of the breakpoint. This property ensures the breakpoint is artificial, and hence the sequence is a circular element. Inversely, this property does not hold when the breakpoint is not artificial (i.e. the sequence is actually an assembly fragment or linear element). Some of these long reads align across the artificial contig breakpoint, indicating these plasmids are extrachromosomal and circular.

### Transfer of predicted plasmid between microbial populations

In duplicate, we streaked *B. fragilis* 214 (donor, carries erythromycin resistant on pFIJ0137_1, one of 14 isolates from Vineis et al. ^55^) and *B. fragilis* 638R (recipient, rifampicin resistant) onto plates with brain-heart infusion agar supplemented with hemin and vitamin K (BHIS). We picked colonies and incubated them in 5 mL BHIS media anaerobically at 37°C for 20 hours. While pFIJ0137_1 lacks conjugation machinery, it contains two relaxases (blue genes in Figure S4) and thus could be mobilized by different conjugative apparatus in the host cell. To mate the donor to the recipient, 250 μL of donor cells were pelleted in a centrifuge at 5,000x gravity. We discarded the supernatant and resuspended the donor in 1 mL of the recipient culture. Again, cells were pelleted at 5,000x gravity, then resuspended in 25 μL of BHIS media. Cells were spotted onto BHIS agar plates and incubated anaerobically for 24 hours. After incubation, cells were resuspended in 1 mL BHIS. 250 μL of this suspension was plated onto BHIS plates containing 8 μg/mL rifampicin and 25 μg/mL erythromycin to select for *B. fragilis* 638R recipients of pFIJ0137_1. Duplicate plates each had approximately 300 colonies. Plating the donor or recipient alone on rifampicin erythromycin plates resulted in zero colonies, confirming the transformants were not spontaneous mutants to either antibiotic. Two transformant colonies were restreaked onto fresh BHIS plates containing 8 ug/mL rifampicin and 25 μg/mL erythromycin. Through short-read sequencing of the donor, recipient, and resulting transconjugants, and by employing a read recruitment analysis, we confirmed that pFIJ0137_1 transferred from B. fragilis 214 to B. fragilis 638R (Figure S4).

### Short-read sequencing of isolate genomes and confirmation of plasmid transfer

We grew 20-hour cultures of *B. fragilis* 214 donor, naive *B. fragilis* 638R, and *B. fragilis* 638R transconjugants containing pFIJ0137_1. Using the QIAseq FX DNA library kit (Qiagen), libraries of these strains were prepared with 100 ng of genomic DNA. DNA was fragmented enzymatically into smaller fragments and desired insert size was achieved by adjusting fragmentation conditions. Fragmented DNA was end repaired and ‘A’s were added to the 3’ ends to stage inserts for ligation. During the ligation step, Illumina compatible Unique Dual Index (UDI) adapters were added to the inserts and the prepared library was PCR amplified. Amplified libraries were cleaned up, and QC was performed using a tapestation. Libraries were sequenced on Illumina MiSeq platform using v2 cassette to generate 2x250bp reads. To confirm the transfer of pFIJ0137_1, we individually recruited reads from the *B. fragilis* 214 donor, naive *B. fragilis* 638R, and *B. fragilis* 638R transconjugants to the pFIJ0137_1 reference sequence. We used anvi’o to create contigs and profile databases (as described above) and visualized these results with the command ‘anvi-interactive’. We independently confirmed the presence of pFIJ0137_1 by assembling genomes using SPAdes^141^ with default parameters.

## Data availability

Reproducible analyses of reference plasmids and chromosomes are available at doi:10.5281/zenodo.5732024. The PlasX model as well as our analyses of known and predicted plasmids are available at doi:10.5281/zenodo.5843600. For all metagenomes, we have compiled the contigs, taxonomic abundances, and PlasX scores at doi:10.5281/zenodo.8175278, gene calls at doi:10.5281/zenodo.5730987, and gene annotations at doi:10.5281/zenodo.5731658. We have deposited long and short sequencing reads from *B. fragilis* isolates into the NCBI Sequence Read Archive (PRJNA782184).

## Code availability

We have released two open-source packages, PlasX (https://github.com/michaelkyu/plasx) and MobMess (https://github.com/michaelkyu/mobmess), along with detailed installation and usage instructions.

## Acknowledgments

We thank Karen Lolans (ORCiD:0000-0003-1903-756X) for performing the long-read sequencing and for providing feedback on the manuscript. We thank Samuel Miller (0000-0002-2836-1401) and Marcus Foo (0000-0003-3436-1632) for their insights into tRNA modification genes. We also thank other members of the Meren Lab at the University of Chicago for their feedback. MKY acknowledges support from Toyota Technological Institute at Chicago.

## Author information

### Contributions

Conceptualization: MKY, AME

Methodology: MKY, ECF, AME

Investigation: MKY, ECF, AME

Visualization: MKY, ECF, AME

Funding acquisition: MKY, AME

Project administration: MKY, AME

Supervision: MKY, AME

Writing – original draft: MKY, ECF

Writing – review & editing: MKY, ECF, AME

Data curation: MKY, ECF

Formal Analysis: MKY, ECF

Resources: ECF, MKY

Software: MKY

Validation: ECF

