## Supplementary Information for "The genetic and ecological landscape of plasmids in the human gut"

[1]Toyota Technological Institute at Chicago; Chicago, IL 60637, USA; [2]Department of Medicine, University of Chicago; Chicago, IL 60637, USA; [3]Graduate Program in the Biological Sciences, University of Chicago; Chicago, IL 60637, USA; [4]Josephine Bay Paul Center for Comparative Molecular Biology and Evolution, Marine Biological Laboratory; Woods Hole, MA 02543, USA; [5]Alfred Wegener Institute for Polar and Marine Research, 27570 Bremerhaven, Germany; [6]Institute for Chemistry and Biology of the Marine Environment, University of Oldenburg, 26129 Oldenburg, Germany; [7]Helmholtz Institute for Functional Marine Biodiversity, 26129 Oldenburg, Germany

### Relevance of PLSDB database for predicting gut plasmids

By training PlasX on all known plasmids in the 2019_03_05 version of the PLSDB database, we designed it to be suitable for identifying plasmids in metagenomes from any environment. The vast majority of plasmids in PLSDB, however, originate from aerobic organisms, with a relatively lesser representation of anaerobic taxa. Our examination of the microbial taxa and isolation sources of the plasmids in PLSDB suggested that the 23% (3772/16168) of plasmids in this database originated from the phylum Bacteroidetes or Firmicutes, each of which includes many organisms that are commonly found in anaerobic human gut environment. In addition, 6% (884) of the plasmids were directly annotated as being isolated from a gut sample. Together, 28% (4536) of plasmids were either isolated from an organism that resolved to Bacteroidetes or Firmicutes, or they were isolated directly from a human gut sample. This unevenness in PLSDB reflects a common limitation of any database, where some types of sequences are more represented than others. To minimize the impact of any single type of sequence from dominating PlasX's logic, and thereby enable it to pay attention to all types of sequences, we explicitly grouped the training sets of plasmid and chromosomal sequences into subtypes and then assigned weights to individual sequences in such a way that every subtype had an equal total weight in the model (Methods). Nevertheless, in theory, the relatively low representation of plasmids from anaerobic organisms in the training dataset can make PlasX more prone to miss true plasmids (“false negatives”) when applied to anaerobic environments such as the human gut microbiome, compared to its applications to environments that are aerobic. Despite the risk of increased proportion of false negatives, PlasX was still able to predict a large number of plasmids from human gut metagenomes. PlasX was also able to classify pWCP as a plasmid (score=0.73), which is a novel plasmid of *Wolbachia* (a likely anaerobe as described by Fallon et al.[1] and Uribe-Alvarez et al.[2]) that was missed by other modern plasmid prediction algorithms.

### Additional validations of predicted plasmids

To determine if a predicted plasmid has canonical plasmid features, we ran MOB-suite[3]. This tool searches a sequence for known examples of four types of features: plasmid replicon (e.g. replication genes), relaxase, mating pair formation, and origin of transfer. We installed MOB-suite v3.0.1 using pip, in an Anaconda Python environment that has mash v2.2. We ran the MOB_typer subroutine (command `mob_typer`) using default parameters and followed the execution instructions at https://github.com/phac-nml/mob-suite, and summarized the results in Table S8.

To determine if a predictive plasmid is a novel sequence, plasmids were blasted against NCBI using the blast package (v2.9.0, installed via bioconda). On October 13, 2021, we downloaded version 5 of the NCBI databases non-redundant nucleotide (nt), ref_prok_rep_genomes, ref_viroids_rep_genomes, and ref_viruses_rep_genomes, and then integrated them into a single database using the `blastdb_aliastool` command. We then searched every predicted plasmids against this combined database, using the `blastn` tool with the ‘-task megablast’ parameter for efficient searching. For each plasmid, we examined all matching NCBI sequences (called 'subjects') and chose the one with the highest ‘qcovs’ (query coverage per subject), which represents the fraction of the plasmid sequence that is covered by all high-scoring segment pairs (HSP). Tiebreaking was done by sorting subjects by the maximum bitscore of the HSPs. If the qcovs of the best matching sequence was ≥90%, then we considered the predicted plasmid as found in NCBI and further categorized the matching sequence by searching for the keywords 'plasmid', 'virus', 'chromosome' (in that order, disregarding capitalization) in its NCBI description. For example, if the description of the matching sequence contained the word 'plasmid', then we said the predicted plasmid matched a known plasmid on NCBI. Similarly, if the description contained 'chromosome' but not 'plasmid' nor 'virus', then we said that the predicted plasmid matched a known chromosome on NCBI. If the qcovs of the best sequence was <90%, then we labeled the predicted plasmid as not found in NCBI.

We further investigated the subset of predictions that were highly similar to a sequence in NCBI and categorized matches as either known plasmids (26.9%), chromosomes (21.3%), viruses (0.6%), or an unclear type of sequence (51.2%) (Figure S2C). A total of 189 predictions matched a known virus. Of these, 110 were recognized as plasmids by MOB-suite or keywords but also contained virus-related COG or Pfam functions, as indicated by the keywords 'virus', 'viral', and 'phage'. These predictions carry both plasmid and viral features, a phenomenon that has previously been reported[4–7]. Surprisingly, 808 predictions that matched a known chromosome were also circular-associated and recognized by MOB-suite or plasmid keywords. One explanation of these data is that these plasmids can switch between an extrachromosomal or a chromosome-integrated state.

We also found that among all of the assembled contigs that were circular, those that were predicted as plasmids tended to have a higher ‘circularity coverage ratio’, defined as the number of reverse-forward read pairs supporting circularity by the read coverage (Figure S2E). Thresholding this ratio could be used as an additional filter in future work to identify plasmids of higher confidence.

### Comparison of PlasX to other tools and new sequences

Here we implemented a more realistic evaluation framework to compare PlasX to three state-of-the-art algorithms, PlasClass[8], PPR-Meta[9], and Platon[10]. We first evaluated performance in 4-fold cross-validation, using a ‘naive’ randomized splitting of sequences into training and test data (Figure S1E). PlasX achieved nearly perfect accuracy, with the highest area under the precision-recall curve (AUCPR=0.99) compared to all other methods (Figure S1F). While naive splitting is a common evaluation technique, it is not a fair strategy as it can separate very similar sequences into training and test data, especially given the redundancy of sequences in public databases, and thus inflate the accuracy of classification. As a more accurate benchmark, we (1) designed an ‘informed’ split by first clustering plasmid and chromosomal sequences into subtypes and then keeping all sequences in the same subtype together in either the training or test data to better evaluate the ability of recognizing novel sequences and (2) assigned normalized weights to sequences to prevent well-studied plasmids from influencing the prediction ability disproportionately (see Methods). This advanced benchmark revealed a greater performance divide between PlasX (weighted AUCPR=0.70) and all other methods, with the next best method performing substantially worse (Platon, weighted AUCPR=0.23) (Figure 1D).

### Additional notes on the execution of other plasmid prediction tools and benchmarks

The purpose of this section is to share installation and runtime details of our benchmarking of PlasX against state-of-the-art tools, Platon[10], PlasClass[8], PPR-Meta[9], and Deeplasmid[11], and our use of the publicly available data for this aim.

We downloaded PlasClass from https://github.com/Shamir-Lab/PlasClass (v0.1.0-2-gb80a4f4). We downloaded PPR-Meta from https://github.com/zhenchengfang/PPR-Meta (v1.0-14-gab99c91). We downloaded Platon from https://github.com/oschwengers/platon, and then modified the code to more efficiently parallelize across many CPUs (modifications at https://github.com/michaelkyu/platon).

To ensure a fair comparison of models in cross-validation (Figures 1D and S1F), we did not use the pretrained versions of PlasClass and Platon from their original studies, but instead we retrained those models using the same training sequences as we used for PlasX in each cross-validation fold. We retrained PlasClass on the 10 kbp slices in each fold. For computational feasibility, we retrained Platon on the whole sequences in each fold, instead of slices. We did not train PlasClass and Platon with sequence weights because they don't take in weights as input, but we did calculate precision and recall with weights. We used Platon's RDS score as its final prediction score, ignoring whether it found other features like conjugation and replication genes, as Platon was unable to identify them in a feasible runtime when evaluating all 10 kbp slices. PPR-Meta and Deeplasmid do not provide software interfaces for retraining new models, so we ran the pretrained versions of these models from their original studies (note that those studies used different sequence datasets).

We downloaded the four sequence versions of the Wolbachia plasmid pWCP from https://doi.org/10.6084/m9.figshare.6380015 (Table S8). We made predictions of pWCP using the original published model versions of PlasClass, Platon, PPR-Meta, and Deeplasmid.

We also downloaded the more recent 2021_06_23_v2 version of PLSDB, which contains 34,513 plasmid sequences. Of these, 21,012 were newly deposited plasmids that were not used to train PlasX, so we focused our evaluation on this subset of plasmids to measure PlasX's generalizability (Table S2). As a comparison, we ran the original pretrained version of Platon in all execution modes ('sensitivity', 'specificity', 'accuracy', and 'characterize').

We downloaded the collection of all ICE sequences (n=552) from ICEberg[12] 2.0 at https://db-mml.sjtu.edu.cn/ICEberg/ on September 30, 2022. We also downloaded 455 prophage sequences from the NCBI Virus data portal (https://www.ncbi.nlm.nih.gov/labs/virus) on September 30, 2022. To download them, we selected the "Bacteriophages" subset from the ">Find Data" menu bar, and then we applied filters of "Only" for the "Provirus" option and "complete" for the "Nucleotide Completeness" option. We made predictions of these ICE's and virus sequences using the original pretrained version of Platon, using its default 'accuracy' mode (Tables S4 and S5).

For the four external datasets described above (pWCP, PLSDB's newer plasmids, ICE's, and viruses), we ran PlasX and other plasmid tools on the whole sequences (instead of 10 kbp slices).

We ran Deeplasmid using the Docker image of the CPU implementation, following instructions at https://github.com/wandreopoulos/deeplasmid (version sha256:10809927e2c8a14cf86231801b804b0bd4bddf600821d17fd8b7e41a15c562c0). While we were able to run Deeplasmid on the Wolbachia plasmid pWCP, it was prohibitively slow to run on the entire set of 10 kbp slices used for cross-validation evaluation. In particular, we found that Deeplasmid running on a MacOS laptop takes ~3 hours for 1,000 slices, so we estimated it would take ~3.7 years to run on all slices. While the GPU implementation of Deeplasmid might be able to run faster, we were unable to execute its prebuilt Docker image (version sha256:f3a22993fb765a7f9678b174245b64976e7e52a4dce85570060900b794af5e43). We suspect that this image is incompatible with modern machine setups, like ours, because Deeplasmid depends on software that is several years old. For example, it requires the CNTK library, for which development was abandoned over 3 years ago (https://docs.microsoft.com/en-us/cognitive-toolkit/releasenotes/cntk_2_7_release_notes). We were also unable to build a new Docker image to run the GPU implementation, despite attempts to modify the Docker build file (see the issue we raised at https://github.com/wandreopoulos/deeplasmid/issues/3).

### Comparison of MobMess to other plasmid clustering methods

To design MobMess, we first examined the histogram of similarities between all pairs of predicted plasmids, revealing an average nucleotide identity (ANI) "valley" with the lowest point at around 85-90% identity (Figure S9A). However, this valley was wide and shallow, reflecting the occurrence of many plasmids that share partial similarity between ~20% to 90% identity. This shallow valley could have emerged due to a number of reasons, such as assemblies from different metagenomes containing sequence fragments that are partially redundant with each other. Another explanation is the dynamism of plasmids—its tendency to recombine with other plasmids, other mobile elements, or chromosomes—resulting in a mosaic composition of genetic material that originated from different sources[13]. In contrast, a deeper valley has been observed when analyzing collections of reference plasmids[14,15] and of bacterial taxa[16,17]. We believe this reflects reference databases containing non-redundant genomes from distant branches of life, such that clear boundaries exist when grouping those genomes into evolutionary clusters. Consequently, previous methods for clustering reference plasmids chose an identity threshold in the middle (e.g. >50% ANI by Redondo-Salvo et al.), as slightly shifting the threshold up or down would have minor effects on clustering. The valley in our collection of metagenome-derived plasmids appeared to lack a clear ANI threshold, as every threshold within the range of 20% to 90% seemed to be almost equally reasonable targets (Figure S9A). Choosing a single threshold appeared to over-split or over-combine plasmids, rather than define ecologically and/or evolutionarily cohesive units. Thus, we designed a new algorithm, MobMess, which does not simply rely on an ANI threshold but, in addition, takes a more nuanced examination of the topology of the sequence similarity network by considering how much one sequence is contained within another sequence.

To identify an appropriate threshold on sequence containment ($I_{local}$ and $C$, defined in Methods), we examined MobMess's behavior across a wide range of thresholds. As the threshold is made stricter, MobMess gradually separated plasmids into distinct clusters, and consequently the number of non-redundant plasmids increased (Figure S9B-C). This growth in non-redundant plasmids occurred at a mostly constant rate from a threshold of 10% to 90%, but it suddenly accelerated from 90% to 100%. These results suggest that a threshold stricter than ≥90% (e.g. ≥95% or ≥99%) would split highly similar plasmids into separate clusters. Thus, we found that ≥90% alignment identity and coverage was a natural threshold to define containment in MobMess.

Other methods have recently been developed to cluster thousands of plasmids[14,15], but unlike MobMess, they are not designed to identify plasmid systems or analyze metagenomic data. To compare methods, we ran MobMess on the same set of 9,894 reference plasmids analyzed by Redondo-Salvo et al[14] (Figure S10). In their study, Redondo-Salvo et al. constructed a plasmid similarity network with 79,727 edges. However, these edges span a wide range of similarity levels, where 66.5% of edges represent an alignment that covers <90% of either sequence (≥10% is not aligned) and 19.0% of edges have <70% alignment coverage (≥30% is not aligned). In contrast, MobMess applies a stricter threshold of ≥90% coverage to construct a smaller but more refined set of 39,680 edges (connecting 25,270 unique pairs of plasmids). Moreover, Redondo-Salvo et al.'s edges are undirected, while MobMess's edges are directed to track smaller versus larger sequences. Retaining this extra information allowed MobMess to distinguish between the 10,860 pairs (43.0%) with unidirectional connections, representing a backbone contained in a compound plasmid, versus the 14,410 pairs (57.0%) with bidirectional connections, representing nearly identical plasmids.

Besides network construction, these methods also diverge in how they conceptually organize plasmids. MobMess dereplicates the 9,894 plasmids into 7,132 non-redundant sequences and then organizes them into 1,044 plasmid systems. In contrast, Redondo-Salvo et al. identified 641 clusters, or 'PTUs'[14]. We found that 135 PTUs did correspond one-to-one to a plasmid system in MobMess, but the other PTUs spanned a wide range of evolutionary relations. At one extreme, 251 PTUs were simple sets of nearly identical plasmids, representing recent and strong relations. At the other extreme, 45 PTUs were complex mixtures of distinct plasmid systems, representing distant and weak relations. For example, the largest PTU contained 2,460 plasmids, which MobMess further dissected into 1,481 non-redundant plasmids and 461 plasmid systems. Figure S11 demonstrates one such plasmid system, where MobMess precisely connects the system's backbone to its compound plasmids in a "star"-like topology, while the approach by Redondo-Salvo et al. connects almost every pair of these plasmids to each other, which obfuscates the internal organization of the plasmid system. Perhaps this is in part because the method by Redondo-Salvo et al. and another related method by Acman et al.[15] have only been tested on reference plasmids that have been completely assembled, while MobMess is designed to handle metagenomic data by distinguishing between fragmented versus complete (circular) plasmids.
