## Supplemental Figures and Tables for "The genetic and ecological landscape of plasmids in the human gut"

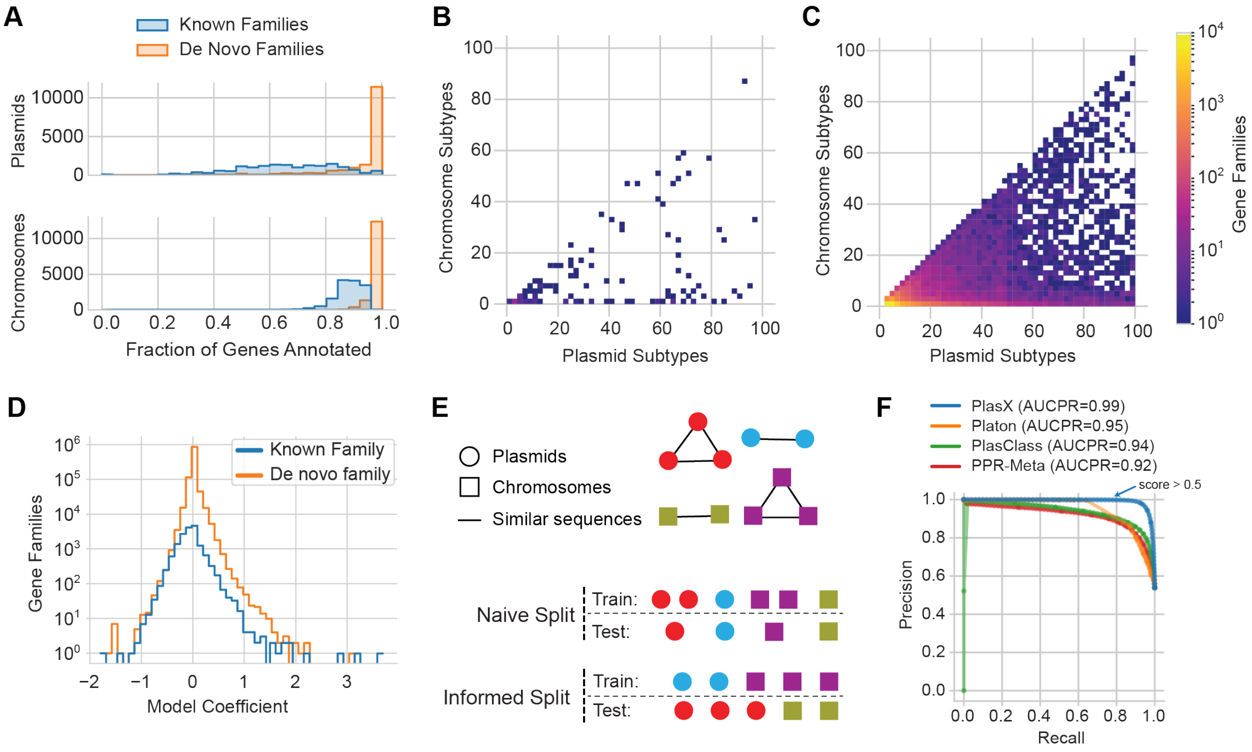


**Figure S1. Additional analysis of PlasX. (A)** Histograms of reference sequences, based on the fraction of genes that have known or *de novo* family annotations. **(B-C)** Two-dimensional histograms of known (B) and *de novo* (C) gene families, based on the number of plasmid and chromosomal subtypes that each family is found in. The number of gene families is log-scaled. Only the gene families that are enriched in plasmid subtypes (i.e. bottom-right triangular region of each plot) are shown. **(D)** Histograms of the coefficients learned by PlasX, showing that the vast majority of coefficients are close to zero. **(E)** Diagrams of different training-test split configurations for cross-validation. A random 'naive' split of plasmids and chromosomal sequences results in training and test sets that have similar sequences, due to the existence of plasmid and chromosomal subtypes that contain highly similar sequences. An 'informed' split assigns all sequences of the same subtype to either training or test, creating a more representative evaluation of a model's ability to generalize to unseen sequences. Colors and edges represent sequences that are in the same subtype. **(F)** Precision-recall curves using 4-fold cross-validation and a naive split.

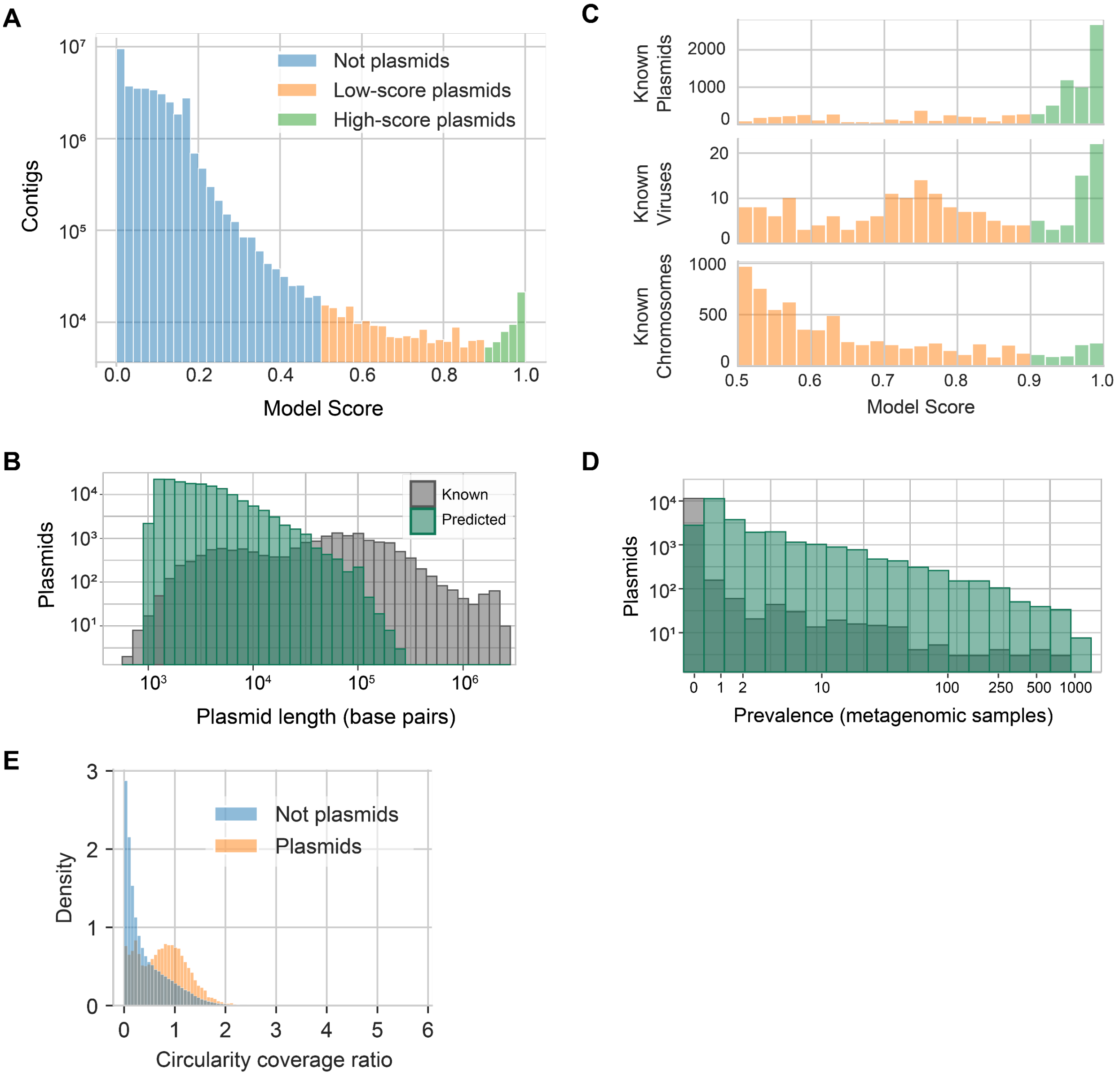


**Figure S2. Additional analysis of predicted plasmids. (A)** Model scores of all contigs assembled from all 1,782 metagenomes. 226,194 plasmids were predicted by applying a score threshold of >0.5. Of these, 50,163 plasmids were high-scoring (≥0.9 score). **(B)** The sequence length of known and predicted plasmids. **(C)** Model scores of predicted plasmids that matched a sequence in NCBI (≥90% alignment identity and ≥90% coverage of the predicted plasmid). Predictions are labeled as a known 'plasmid', 'virus', or 'chromosome' based on the presence of these words in the description of the matching NCBI sequence. We searched NCBI for only the filtered set of 100,719 non-fragment predictions. **(D)** The prevalence of reference and predicted plasmids across all metagenomes. **(E)** We calculated a “circularity coverage ratio” as the number of supporting reverse-forward reads divided by the average coverage of a contig. All circular contigs are shown, and they are colored if they were predicted by PlasX as plasmids with score >0.5 (orange) or not plasmids (blue).

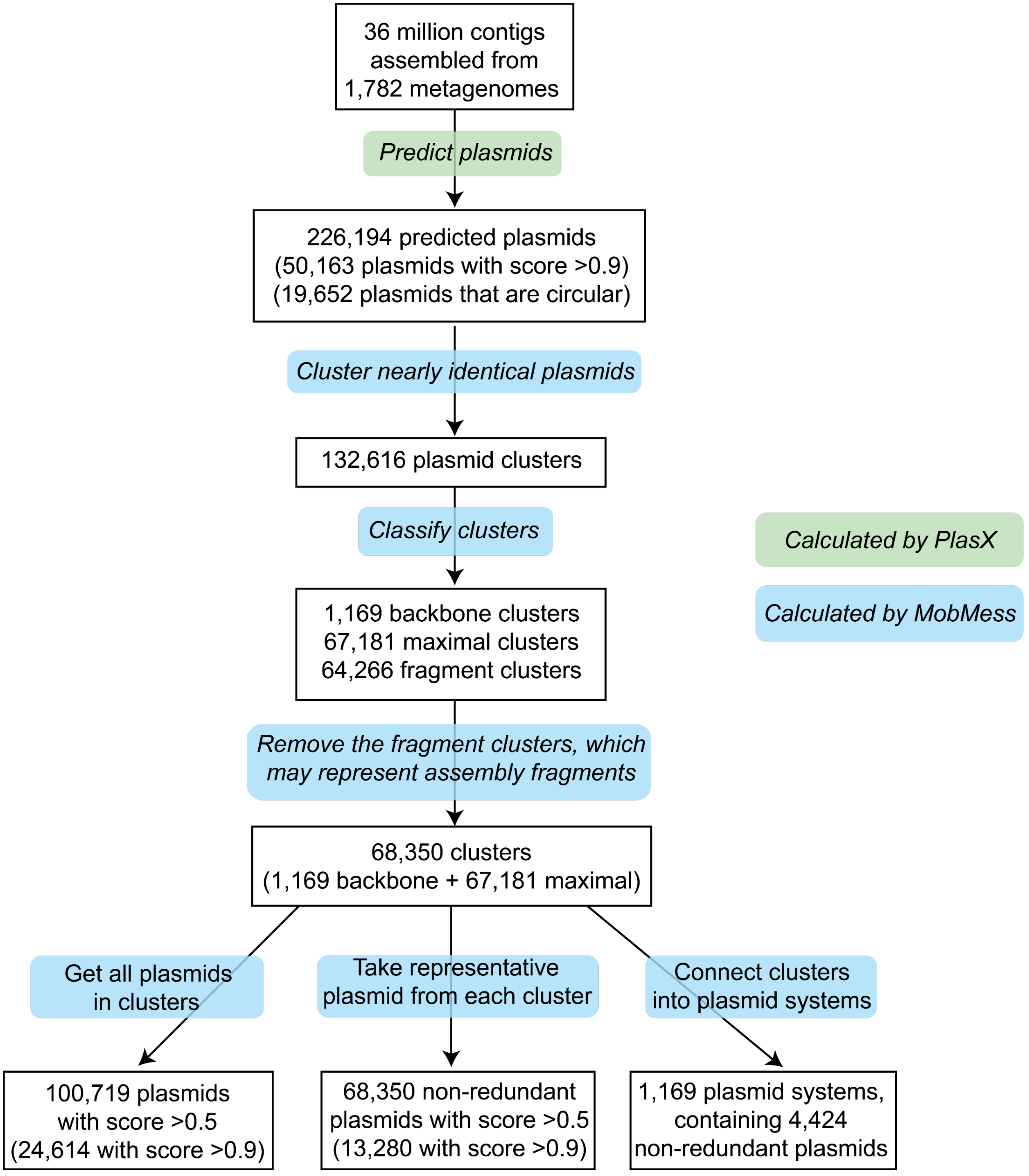


**Figure S3. Workflow of predicting plasmids with PlasX and organizing them with MobMess.**

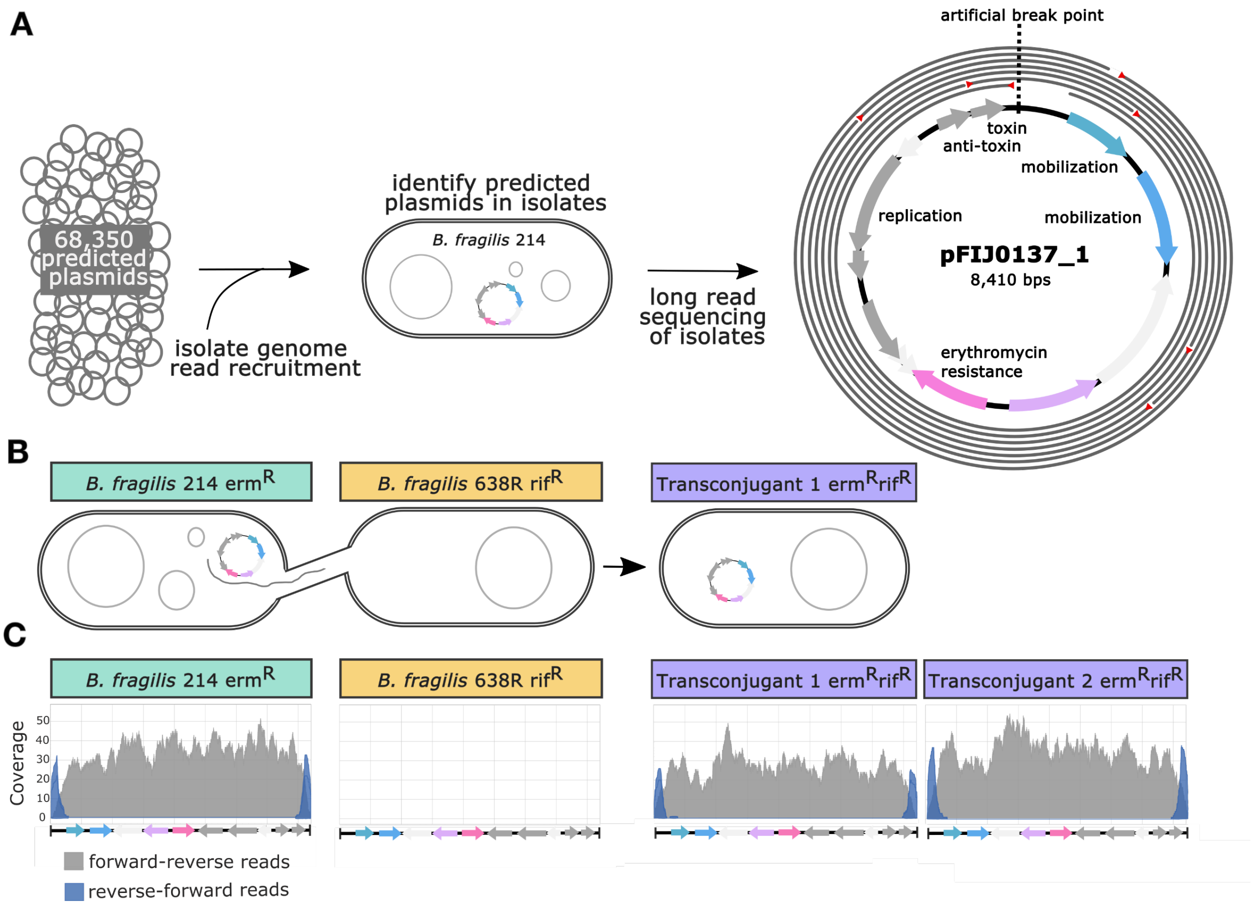


**Figure S4. Experimental validation of plasmid predictions. (A)** We recruited reads from the sequenced genomes of 14 *Bacteroides* isolates to determine which isolates contain our predicted plasmids. We further confirmed the presence and circularity of a predicted plasmid, pFIJ1037_1, in the isolate *B. fragilis* 214 by long read sequencing. Gray circles represent 7 (of 500) long reads that align to pFIJ0137_1. Red triangles designate the beginning of a long read. **(B)** Transfer of pFIJ1037_1 from *B. fragilis* 214 to *B. fragilis* 638R via conjugation and selection on erythromycin- and rifampicin-containing media. **(C)** Coverage plots showing read recruitment of *B. fragilis* whole-genome sequencing reads to the pFIJ1037_1 reference sequence, confirming transfer of pFIJ1037_1. Gray are forward-reverse reads, while blue are reverse-forward reads that indicate the circularity of pFIJ1037_1.

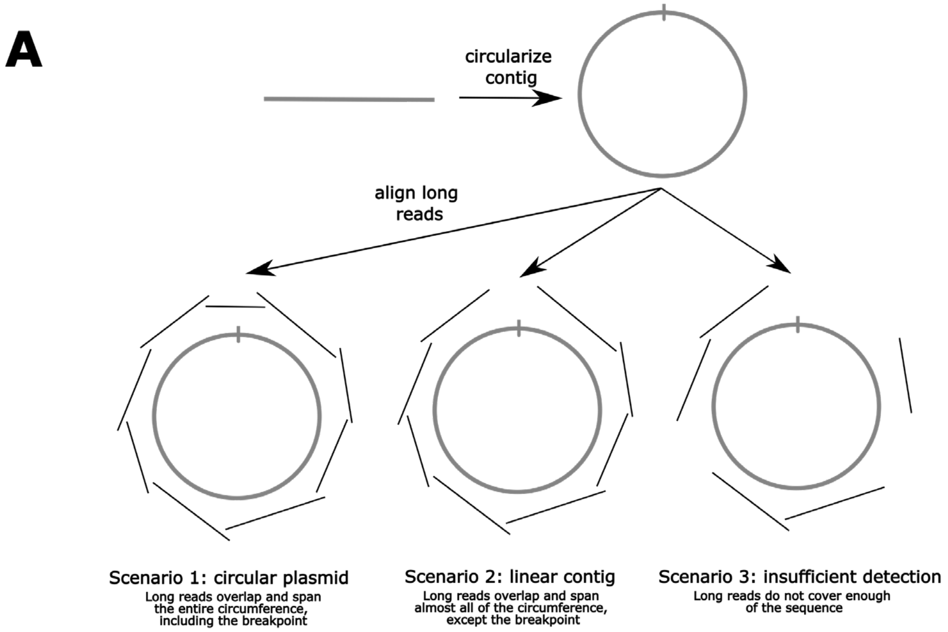


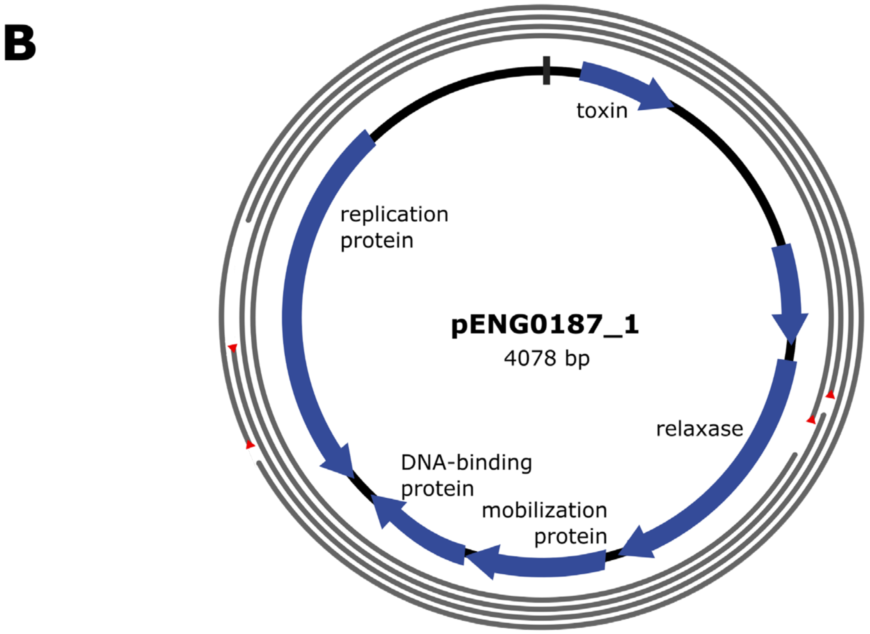


**Figure S5. Long read circularity. (A)** The process to identify circular plasmids using long read sequences. Contigs are always assembled as linear sequences even when originally circular in the environment. We can determine their original configuration by aligning long reads around the entire sequence. **(B)** Long reads from *B. fragilis* 216 aligned to pENG0187_1, demonstrating circularity. 4 of 500 reads are shown for simplicity. Red triangles designate the beginning of a long read.

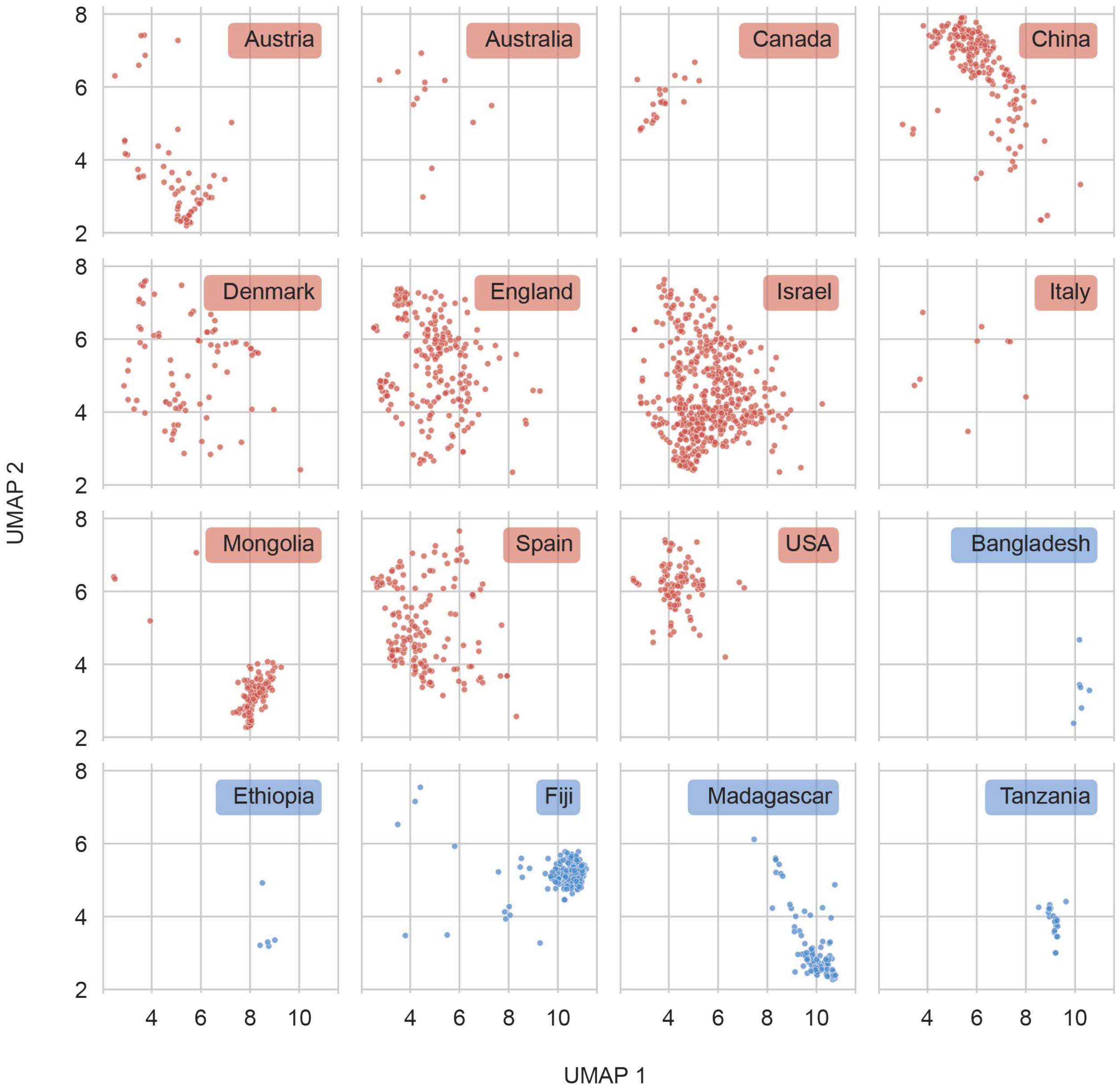


**Figure S6. UMAP plot as in Figure 3C.** Metagenomes have been partitioned to show clustering within each country.

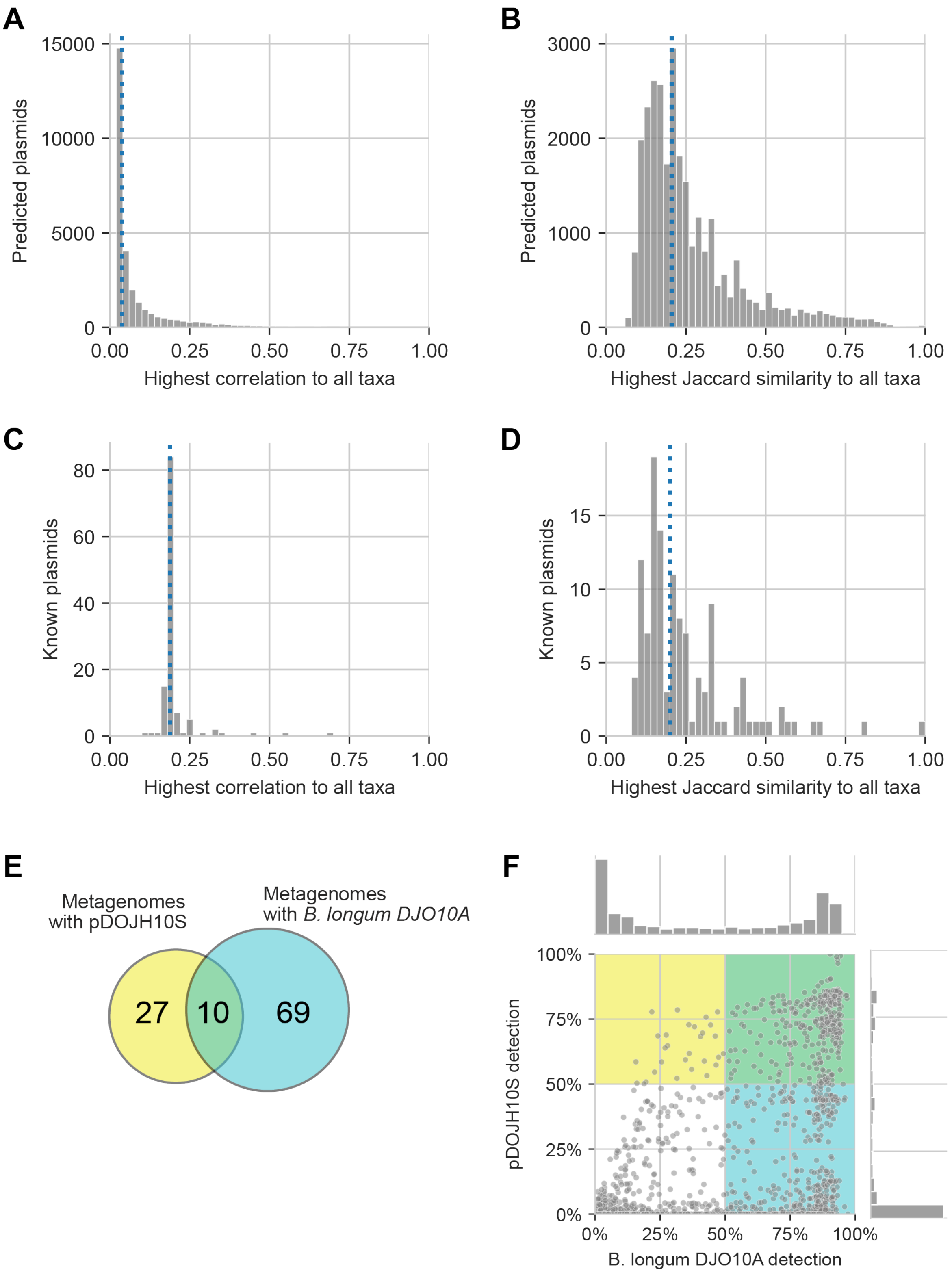


**Figure S7. Comparison of the ecological distributions of plasmids and microbial taxonomy.** We measured the association between every plasmid and taxon by calculating the correlation between their abundance levels across metagenomes, using the SparCC technique[66]. As another association measure, we applied thresholds to the abundance levels and then calculated the Jaccard similarity between the metagenomes containing the plasmid versus those containing the taxon. We estimated taxon abundances with bracken[67]. We restricted analyses to plasmids that were present in at least 5 metagenomes. **(A-B)** For every predicted plasmid, we identified the taxon with the highest correlation **(A)** or Jaccard similarity **(B)**. **(C-D)** We did the same to identify the best matching taxa of reference plasmids. Blue lines indicate the median of each distribution. **(E)** Venn diagram showing the discordance between the metagenomes containing a plasmid pDOJH10S and those containing its cognate host, a *B. longum* strain. **(F)** Detection of pDOJH10S and the *B. longum* strain, based on read recruitment instead of bracken. Each point represents a metagenome. Yellow and blue rectangles highlight the metagenomes where the plasmid or strain, respectively, are identified as being present with >50% detection.

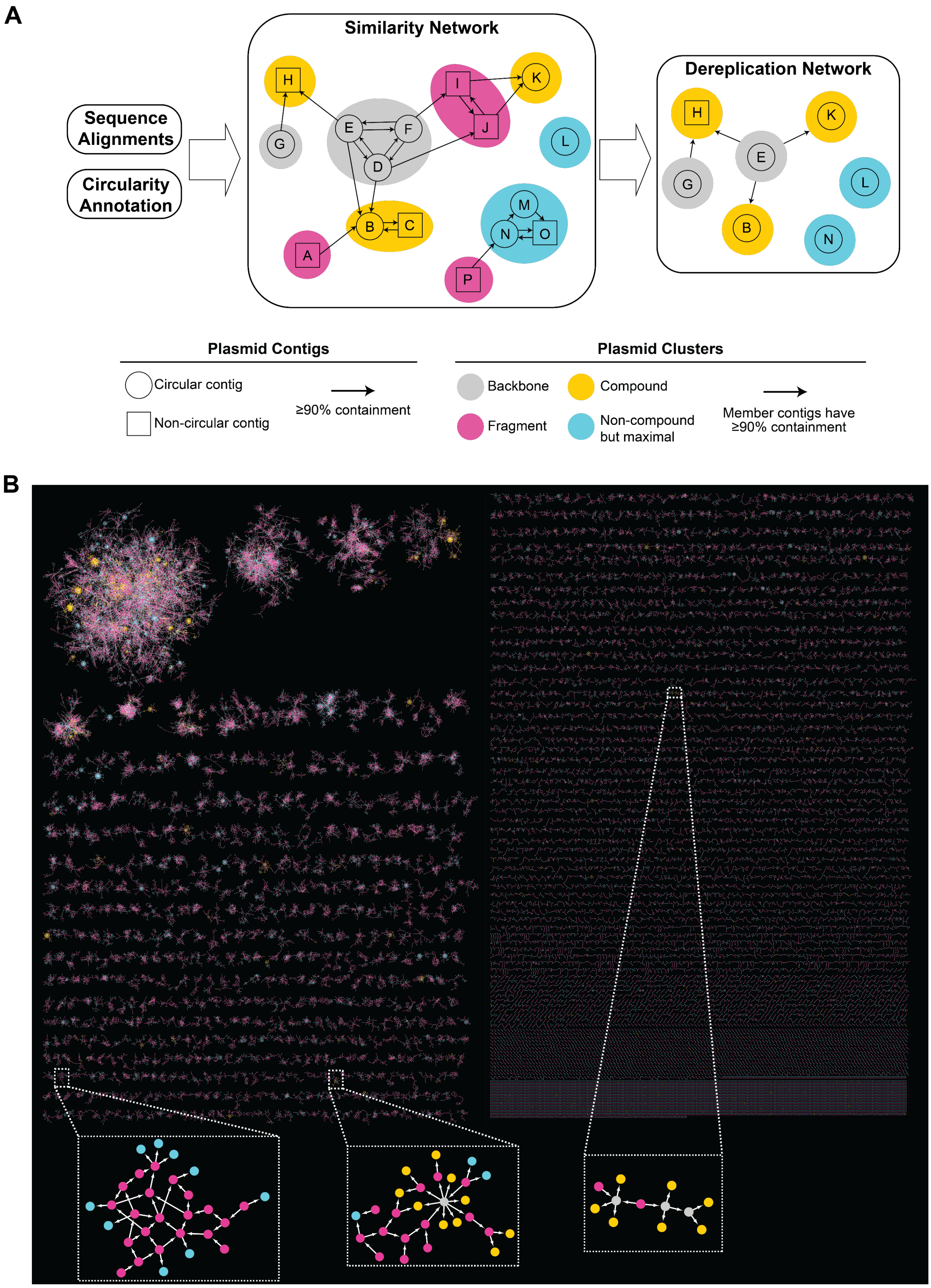
A
Sequence Alignments
Circularity Annotation
Similarity Network
Dereplication Network
Plasmid Contigs
Circular contig
Non-circular contig
≥90% containment
Plasmid Clusters
Backbone
Fragment
Compound
Non-compound but maximal
Member contigs have ≥90% containment
B

**Figure S8. The MobMess algorithm and application to predicted plasmids. (A)** Diagram of the MobMess algorithm for dereplicating plasmids and discovering plasmid systems. All-vs-all sequence alignments and circularity information are used to construct a similarity network of plasmid contigs. Similar contigs are clustered, and every cluster is labeled as either a backbone, fragment, compound, or non-compound maximal. A plasmid system consists of a backbone cluster and the compound clusters connected to the backbone. This example shows two systems: one system has G as the backbone (H is the compound plasmid), and another system has D, E, and F as the backbone (B, C, H, and K are the compound plasmids). To dereplicate, fragment clusters are discarded and a representative sequence is chosen for every non-fragment cluster. **(B)** Network of clusters of predicted plasmids. All clusters are shown except those that are not connected to any other cluster.

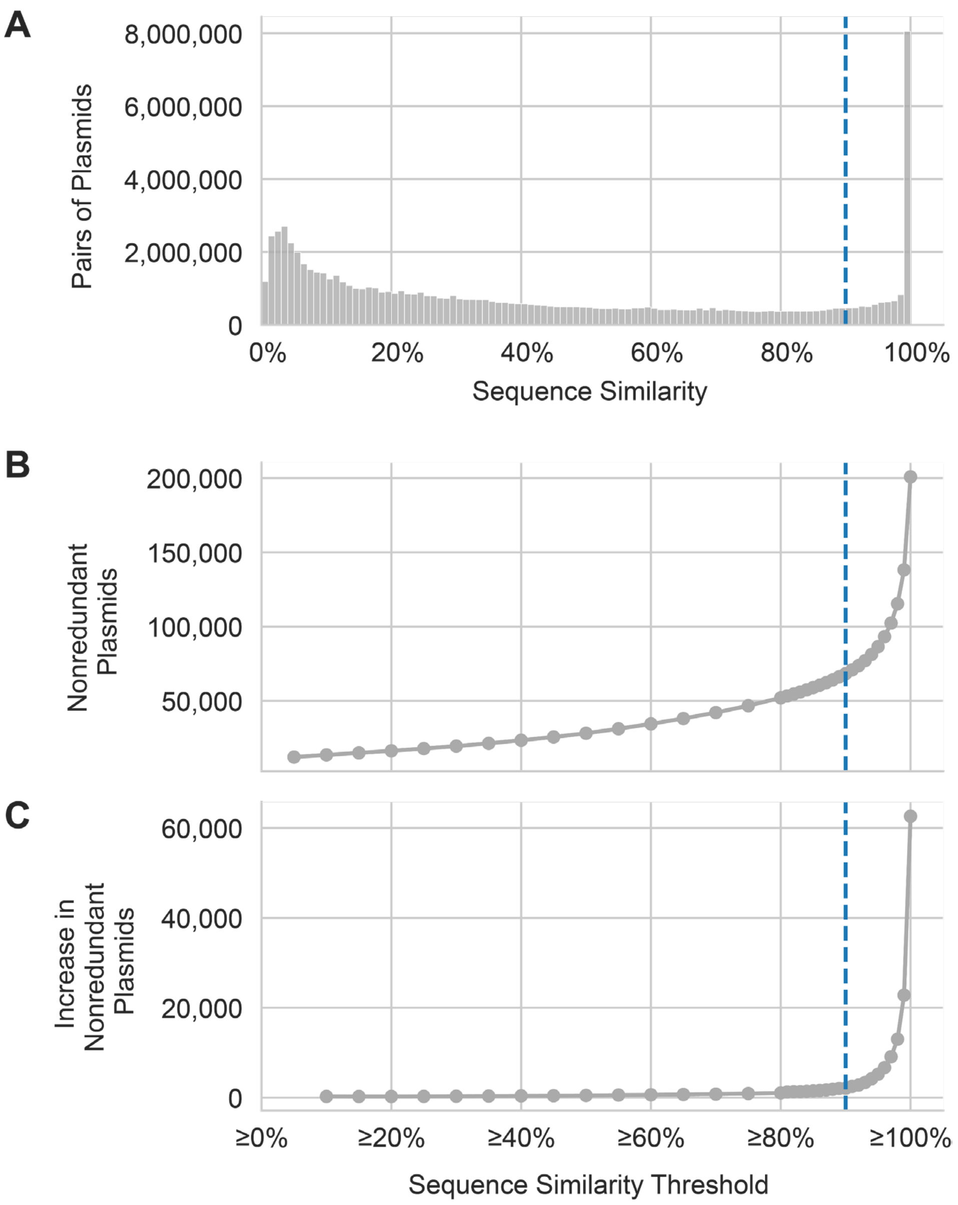


**Figure S9. Choosing a similarity threshold for MobMess. (A)** Histogram of similarities between every pair of the 226,194 predicted plasmid contigs. **(B)** We ran MobMess using different thresholds on the similarity, and then we calculated the number of non-redundant plasmids generated. **(C)** The derivative of the curve in B. The blue dashed lines represent our current ≥90% similarity threshold.

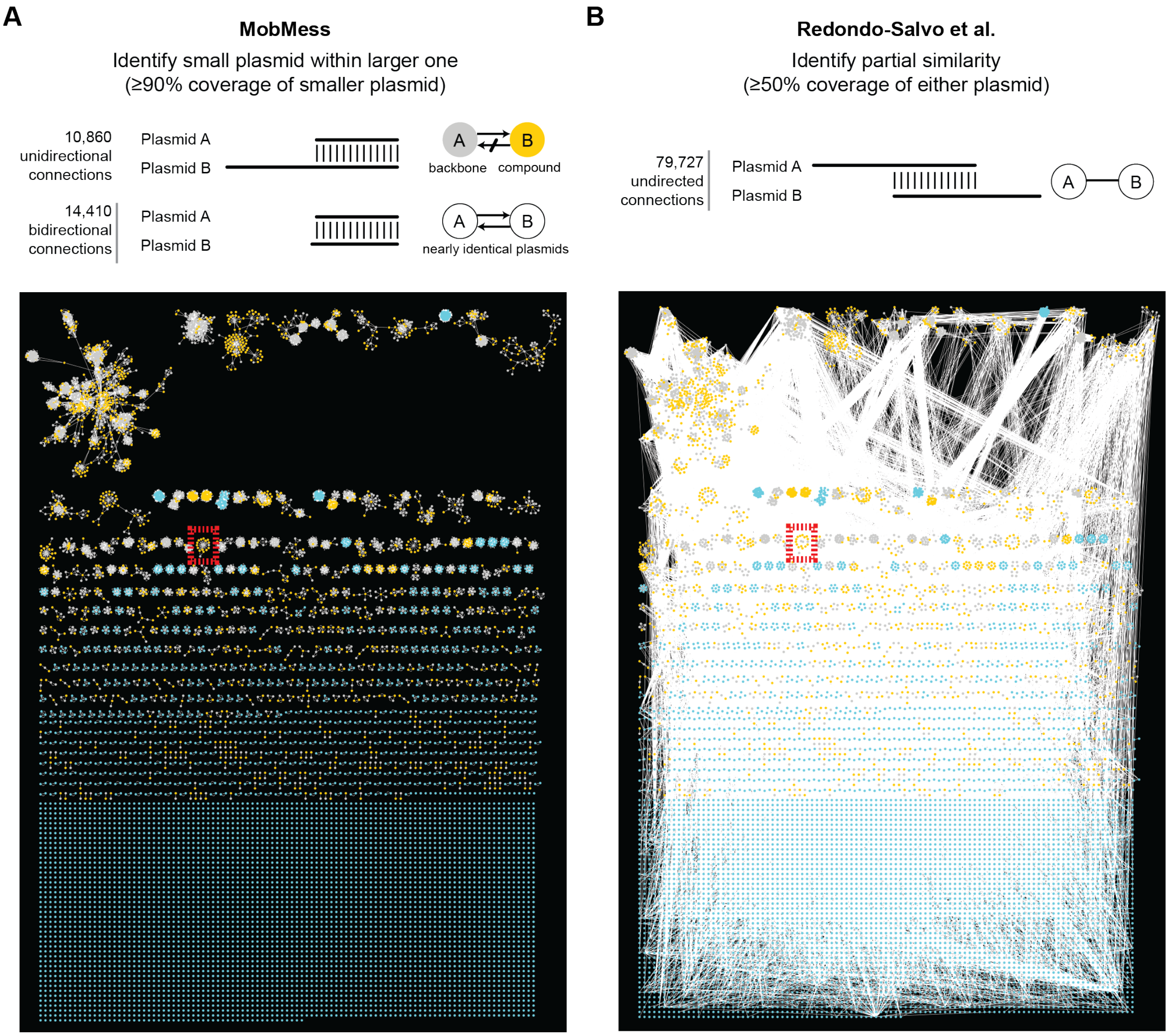


**Figure S10. Conceptual differences in constructing plasmid similarity networks.** We ran MobMess on the set of 9,894 reference plasmids analyzed by Redondo-Salvo et al.[72]. MobMess constructs a network with directed edges, by aligning plasmids and determining if one plasmid is found as a subsequence within another. Redondo-Salvo et al. constructs a network with undirected edges, by determining whether two plasmids contain partial homology. **(A-B)** Visualization of the similarity networks. We used Cytoscape[74] and the Prefuse directed layout algorithm[75] to lay out the nodes in the MobMess network (A), and then we applied the same layout to the Redondo-Salvo et al. network (B). The red boxes represent the example shown in Figure S11.

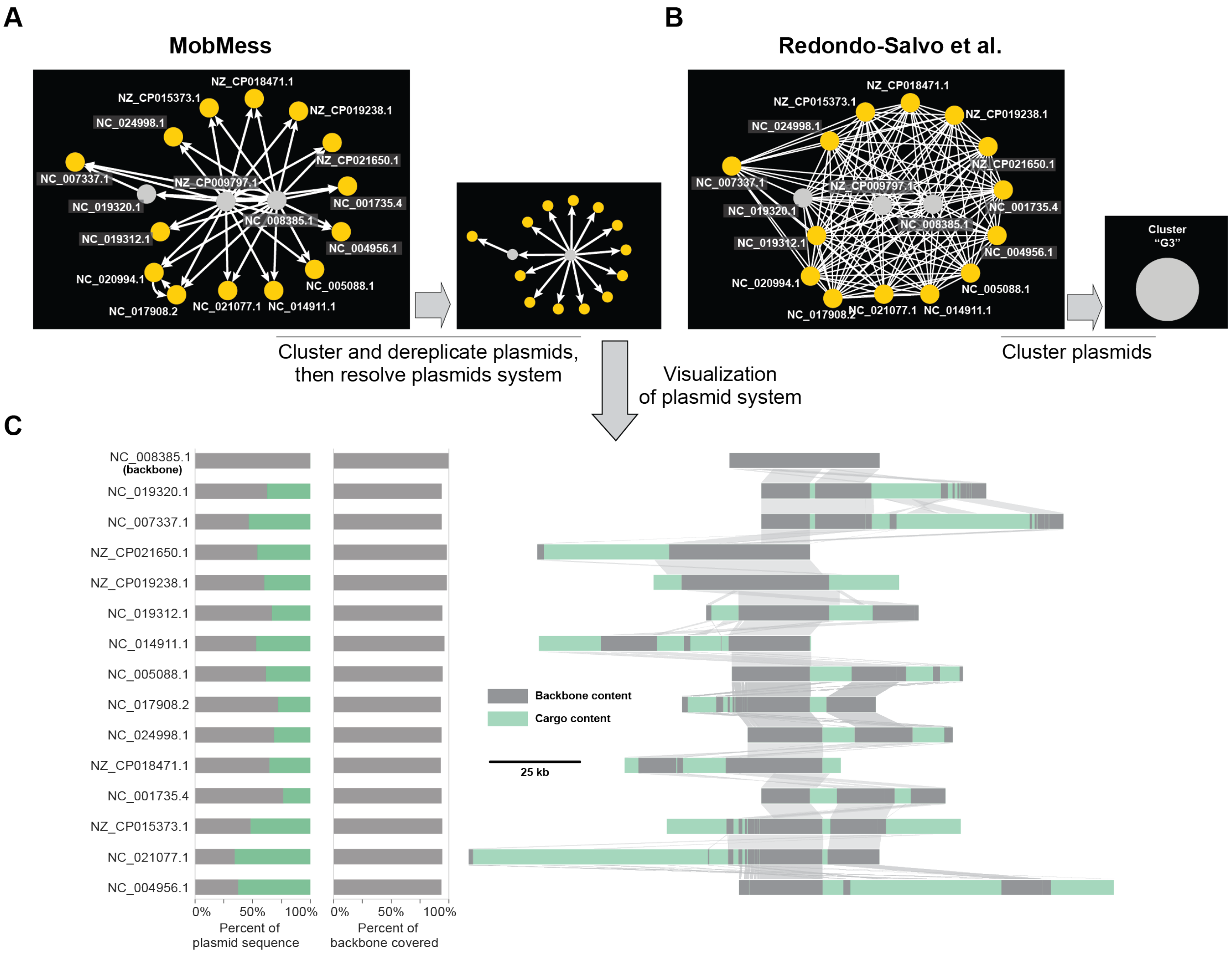


**Figure S11. Comparison of MobMess versus Redondo-Salvo et al.[72] for studying a plasmid system. (A-B)** An example from the similarity networks in Figure S10, showing the connections between 17 plasmids from the same plasmid system. MobMess further collapses its network to dereplicate plasmids and reveal the plasmid systems's "star"-like topology, where a backbone connects to its compound plasmids. Redondo-Salvo et al. did recognize that these plasmids are related (represented by a cluster called "G3"), but they connected almost every pair of these plasmids in a "hairball" topology, obfuscating the system's internal organization. **(C)** Alignments of plasmids in the MobMess system. Subregions in every sequence are colored gray or green to represent backbone or cargo content, respectively. Ribbons between sequences represent the alignment of subregions. The barcharts show the total breakdown of each plasmid into backbone versus cargo, as well as the fraction of the backbone sequence ('NC_008385.1') that is found within the plasmid.

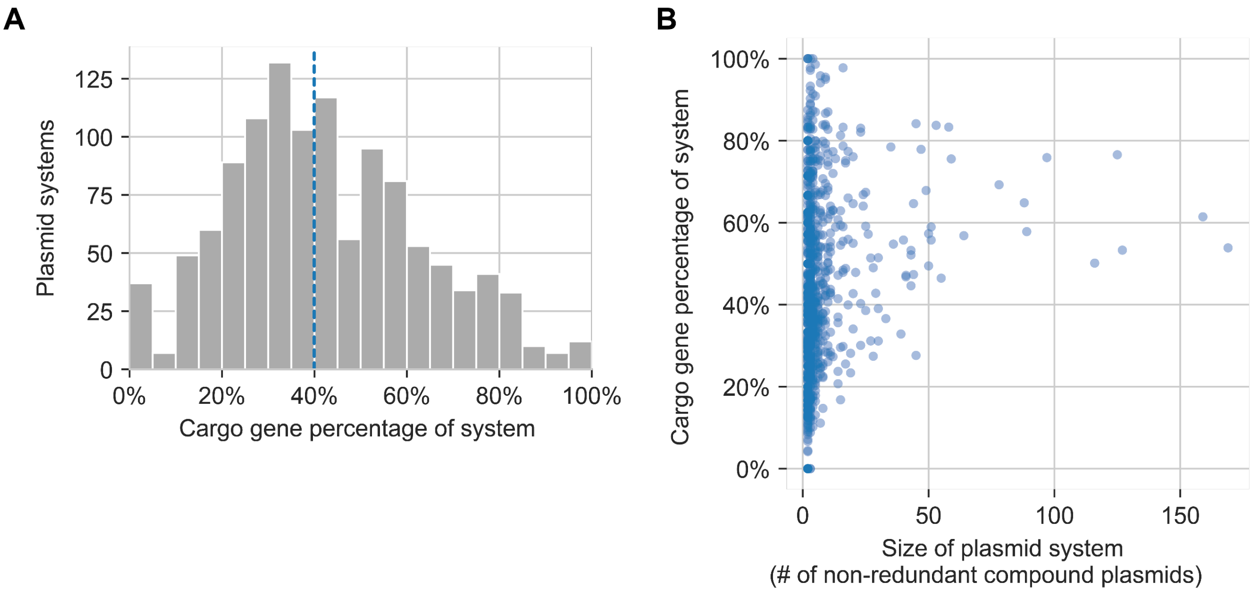


**Figure S12. Backbone and cargo composition of plasmid systems. (A)** For every plasmid system and compound plasmid in the system, we calculated the percentage of genes on the compound plasmid that were classified as cargo versus backbone genes (see Methods). We then averaged the cargo gene percentages across all compound plasmids in the system (x-axis). The vertical blue line shows the median at 40%. **(B)** Scatterplot of the cargo gene percentage versus the size of a plasmid system, showing a lack of correlation ($R^2$ = 0.03). We defined the size as the number of non-redundant compound plasmids.

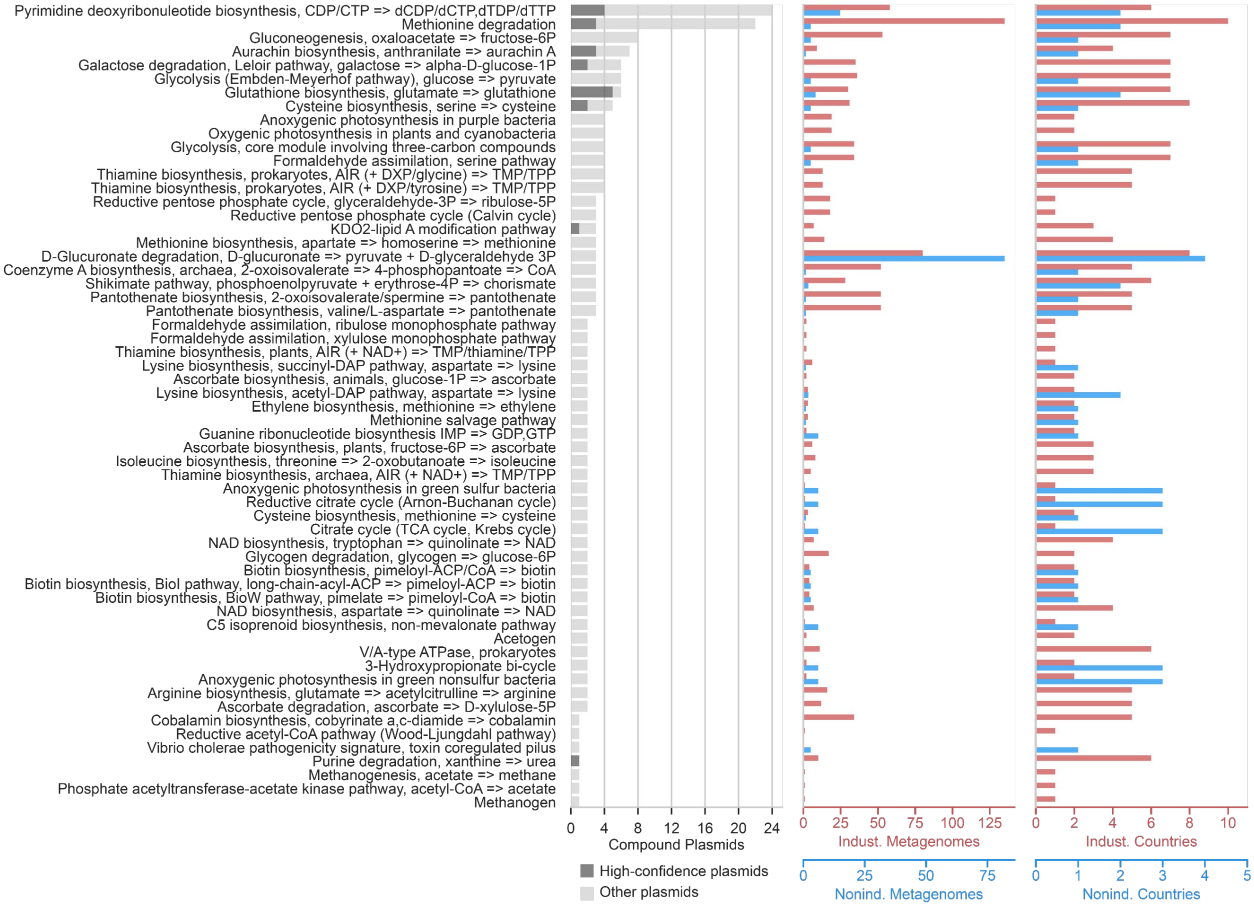


**Figure S13. Functional annotation of cargo genes to KEGG modules, similar to Figures 5A and 5B.** This plot excludes KEGG modules that occur in only one plasmid system. To avoid redundancy with Figure 5A, this plot also excludes modules that occur in cargo genes annotated to antibiotic resistance.

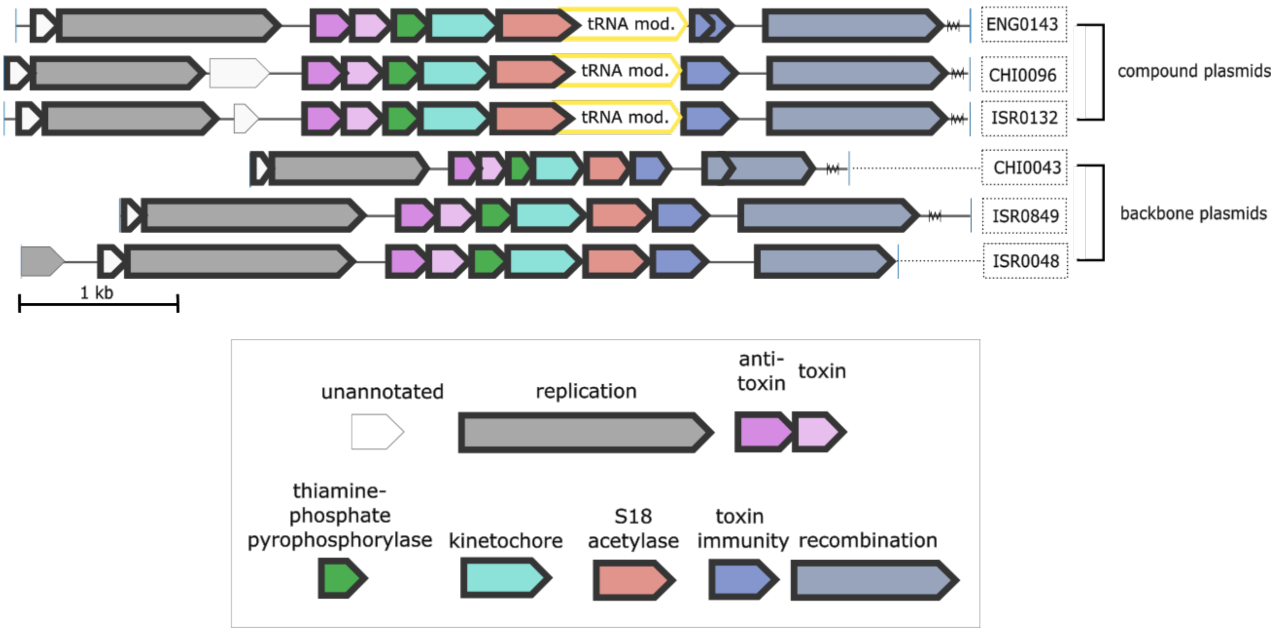


**Figure S14. Plasmid system PS1110**. Compound plasmids in this system contain a gene that encodes two enzymes, a tRNA Gm18 2'-O-methylase (yellow, 'tRNA mod.') and a Ribosomal protein S18 acetylase (red). Backbone plasmids contain a similar gene that encodes the S18 acetylase but lacks the tRNA methylase. Backbone genes have a thick, black outline.

### Supplemental Tables

**Table S1.**

Summary of reference plasmids and chromosomes

**Table S2.**

Prediction of plasmids in the latest version of PLSDB (2020_06_23_v2) by PlasX and Platon

**Table S3.**

Prediction of a Wolbachia plasmid

**Table S4.**

Prediction of ICEs as plasmids by PlasX and Platon

**Table S5.**

Prediction of prophages as plasmids by PlasX and Platon

**Table S6.**

COGs and Pfams ranked by their PlasX coefficients

**Table S7.**

Names, accession numbers, and metadata of metagenomes

**Table S8.**

Summary of predicted plasmids (model scores, orthogonal support, circularity, and NCBI blast results)

**Table S9.**

Summary of plasmid systems

**Table S10.**

The length and percent circularity of plasmids that are part of a system versus plasmids that are *not* part of any system.

**Table S11.**

Gene sequences for plasmid systems shown in Figure 4D, Figure S14 and Figure 5F

**Table S12.**

DNA extraction and sequencing parameters for long read sequencing of isolate genomes

**Table S13.**

Read recruitment detection of pDOJH10S and *B. longum DJO10A*
